# Heat shock induces silent ribosomes and reorganizes mRNA turnover

**DOI:** 10.1101/2025.09.26.678833

**Authors:** Sayanur Rahaman, Nicole Schiffelholz, Nitish Mittal, Klemens E. Fröhlich, Mihaela Zavolan, Attila Becskei

## Abstract

mRNAs associate with single or multiple ribosomes; these ribosomal assemblies — monosomes and polysomes — translate the mRNAs before degradation. The impact of heat stress on this mRNA turnover remains unclear. In heat-shocked yeast cells, the proportion of monosomes within the ribosomal assemblies rises without a corresponding increase in the number of mRNAs associated with them. As a result, most monosomes are devoid of mRNAs and silent, lacking translational initiation factors and proteins facilitating posttranslational folding. The accumulation of silent ribosomes generally reduces the rate of association of transcripts with the ribosomes. However, elevated temperatures enhance the ribosomal association of specific mRNAs, primarily those encoding heat-shock proteins, allowing them to balance their increased degradation rates. Additionally, reduced binding of the Xrn1 exonuclease to mRNAs diminishes the influence of codon optimality on mRNA stability. These mechanisms reorganize mRNA turnover to prioritize heat-shock protein synthesis over ribosome biogenesis.

## INTRODUCTION

Upon being exported to the cytoplasm, mRNAs associate with the small subunit of the ribosome to initiate translation, following which the full ribosome is assembled^1,2^. At a given time, one or more ribosomes are associated with an mRNA; these monosomes and polysomes carry out the translation of mRNAs^3,4^. After multiple rounds of translation, most mRNA molecules are degraded co-translationally, completing the last step of the cytoplasmic turnover of mRNAs^5^. The bulk of the mRNAs is degraded in the 5’-to-3’ direction by Xrn1, while the degradation in the 3’-to-5’ direction occurs through the exosomes^6,7^.

Translation and mRNA degradation are closely interconnected. mRNAs enriched in optimal codons are translated more efficiently and exhibit prolonged half-lives^8^. Conversely, the disruption of translation by mutating the translation initiation codon abolishes the differences in mRNA stability^9^, underscoring the link between RNA translation and degradation. Since optimal codons are recognized faster by their highly-abundant cognate tRNAs, the resulting faster elongation is thought to stabilize mRNAs^8^. Similar mechanisms underlie mRNA degradation triggered by quality control pathways^10^.

mRNAs exist in different complexation states—free, monosome-associated, and polysome-associated. Their distribution among these states is influenced by multiple factors. For example, mRNAs responsive to codon optimality are longer and mainly polysome-associated. In contrast, monosome-associated mRNAs do not exhibit this behavior: their translation efficiencies and half-lives remain unaffected by codon optimality although their translation efficiency can surpass that of the longer polysome-associated mRNAs^11^. Other phenomena can also affect the distribution: inhibition of translation initiation and elongation are expected to shift mRNAs to free and monosome-associated forms, respectively.

Translation elongation can be affected by environmental stress, such as heat shock^12^, which can alter the proportion of mRNA complexation states. Furthermore, heat shock induces the expression of chaperones and heat shock proteins but suppresses that of mRNAs encoding ribosomal proteins^13^. Consequently, the assembly of ribosomes is reduced, particularly upon severe heat stress^14^. Since heat shock affects the expression of the components of the translational apparatus as well as the processes influencing the proportion of complexation states of the mRNA, it is unclear how elevated temperatures affect the balance between association and degradation steps during mRNA turnover.

We aimed to investigate how heat shock alters the complexation states and turnover of mRNA. As high temperatures complicate the measurement of reaction rates, including RNA half-lives^15^, we optimized metabolic labeling under heat shock conditions and compared changes in degradation rates of mRNAs with the rates of their ribosomal association.

## RESULTS

### Contrasting Responses of the Ribosomes and mRNAs in the Ribosome Assemblies to Heat Shock

To investigate the response of the ribosomal assemblies to heat shock, we compared the polysome profiles of yeast cells grown at 30°C with those measured 30 minutes after shifting the cells to 42°C (Fig. 1A). Heat shock induced a significant restructuring of the polysome profile, characterized by a pronounced increase in the monosome peak and a concurrent reduction in polysome fractions.

**Figure 1.**
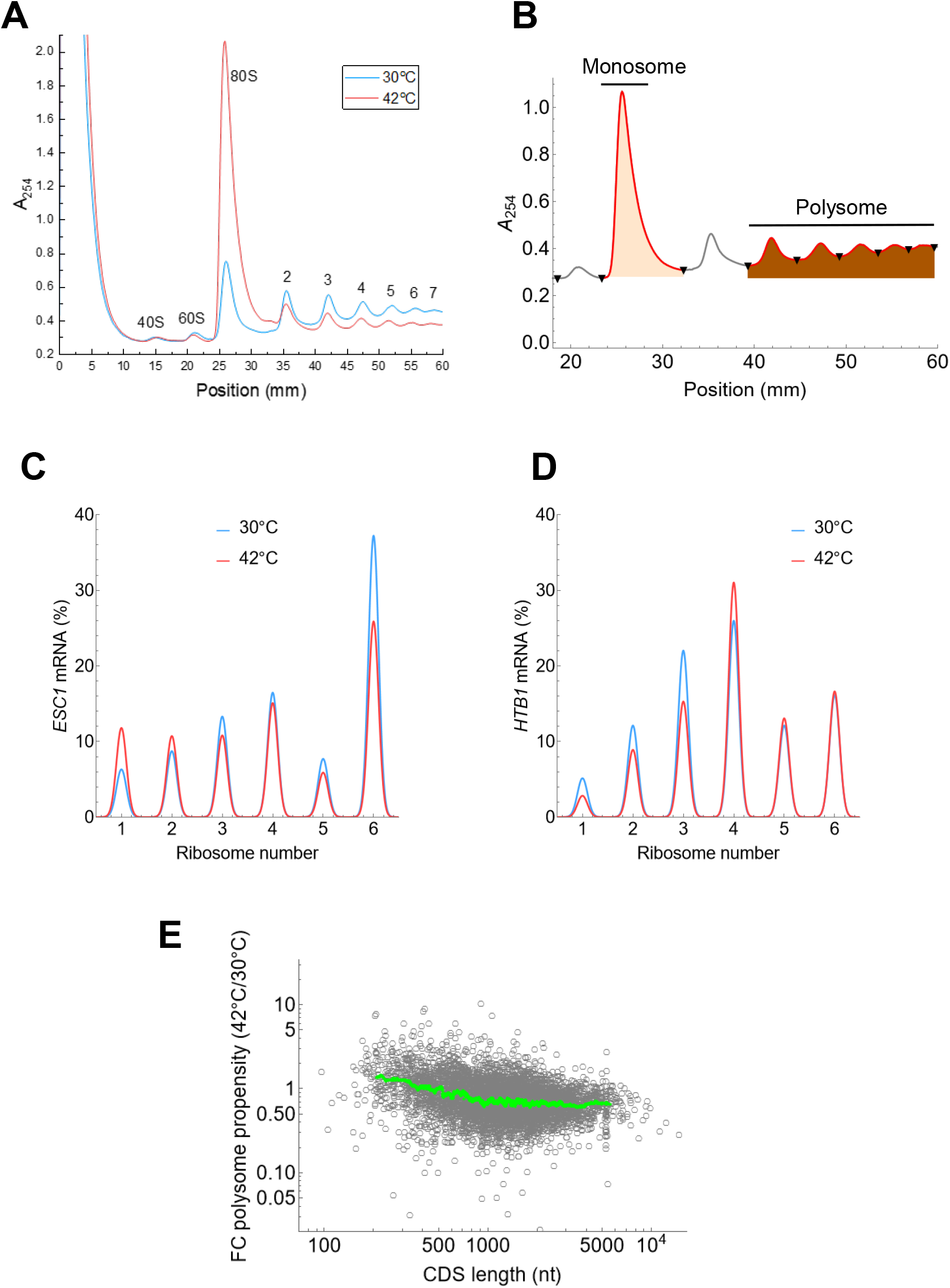
Polysome profiling of cells exposed to heat shock. **(A)** The ribosome counts in the polysome profiles are indicated at the absorbance peaks (A_254_) relative to their positions (mm) in the ultracentrifuge tube. Yeast cells were incubated at 30°C or heat shocked at 42°C for 30 min. **(B)** The shaded areas represent the total absorbance in the monosome (57.1) and polysome (three and more ribosomes: 20.3) peaks. The baseline is defined by the lowest absorbance in the polysome profile. **(C)** The peaks of the bell-shaped curves represent the frequency (in percentages) of the *ESC1* mRNA distributed across different mRNA complexation states, as measured by RT-qPCR. The sum of all peaks, including the free, 40S, 60S and ribosome bound fractions, totals 100%. The ribosome number = 6 includes mRNAs associated with 6 and more ribosomes. *MNR* = 4.3 and 3.4, *PP* = 11.8 and 4.9 at 30°C and 42°C, respectively. **(D)** The distribution of the *HTB1* mRNA across the fractions (as in C). MNR = 3.7 and 3.7, PP = 14.8 and 26.9 at 30°C and 42°C, respectively. **(E)** The fold-change in the polysome propensity of mRNAs measured at 30°C and 42°C. The green line is the moving average.

To quantify these changes more precisely, we calculated the polysome propensity, defined as the ratio of a parameter measured in fractions containing three or more ribosomes to that in the monosome fractions (Fig. 1B). Using absorbance as the measured parameter, the distribution of the ribosomes in the transcriptome can be assessed, allowing the calculation of the ribosomal polysome propensity. This ribosomal polysome propensity decreased substantially upon heat shock, dropping nearly sevenfold from 2.8 to 0.4 (Fig. 1A, B). This decline underscores a dramatic shift in the ribosomal assemblies following exposure to elevated temperature.

We next examined the distribution of mRNA among the ribosomal assemblies by analyzing two representative transcripts: a short mRNA, *HTB1* (396 nucleotides-long coding sequence (CDS)), and a long mRNA, *ESC1* (4977 nucleotides-long CDS). mRNA relative abundances were used as the measured parameter to calculate the mRNA polysome propensities. For *ESC1*, there was a slight increase in the monosome fraction upon heat shock, resulting in a twofold decrease in the mRNA polysome propensity (Fig. 1C). In contrast, *HTB1* exhibited a reduction in monosome association, leading to a twofold increase in the mRNA polysome propensity (Fig. 1D). Thus, while the absorbance profiles showed marked shifts, changes in the distribution of selected mRNAs were comparatively minor and exhibited transcript-specific variations.

To evaluate polysome propensity of the entire transcriptome, we performed RNA-seq on polysome fractions. The median mRNA polysome propensity at 42°C was 4.3 (Data S1), which is only slightly lower than that measured at 30°C^11^, with the median mRNA polysome propensity decreasing by only 0.72-fold. mRNAs with shorter coding sequences changed even less or not at all (Fig. 1E).

What explains the inconsistency between the polysome propensity of the ribosomes and mRNAs? We hypothesize that the prominent increase in the monosome peak, despite the minimal change in the mRNA polysome propensity of the transcriptome, might be due to the presence of ribosomes devoid of mRNAs.

### Most monosomes are devoid of mRNAs upon heat shock

To test the hypothesis that monosomes might lack mRNAs, we compared the TPMs of total mRNA (∑mRNA) and ribosomal RNA (rRNA) in each fraction. We performed RNA-seq on total RNA samples that had not undergone rRNA depletion, allowing us to detect all rRNA species. Most rRNA species were proportionally represented across the fractions, except for 5S rRNA. The 18S/25S rRNA and 5.8S/25S rRNA proportions showed minimal or less than 2fold variations across the fractions (Fig. S1A-B). The 5S rRNA showed an anomalous amplification of RNA-seq counts in the monosome fraction (Fig. S1C), despite displaying proportional intensities when measured by polyacrylamide gel electrophoresis (PAGE) (Fig. S2A). Therefore, we calculated the ∑mRNA/∑rRNA ratios in which the total rRNAs excluded the 5S rRNA.

The ∑mRNA/∑rRNA ratios in the tetrasome pools were nearly identical at 30°C and 42°C. In contrast, the monosome fraction showed significant differences (Fig. 2A). Assuming all monosomes are fully loaded with mRNA (100%) at 30°C, these two ∑mRNA/∑rRNA ratios suggest that only 20.8% of monosomes are associated with mRNA at 42°C.

**Figure 2.**
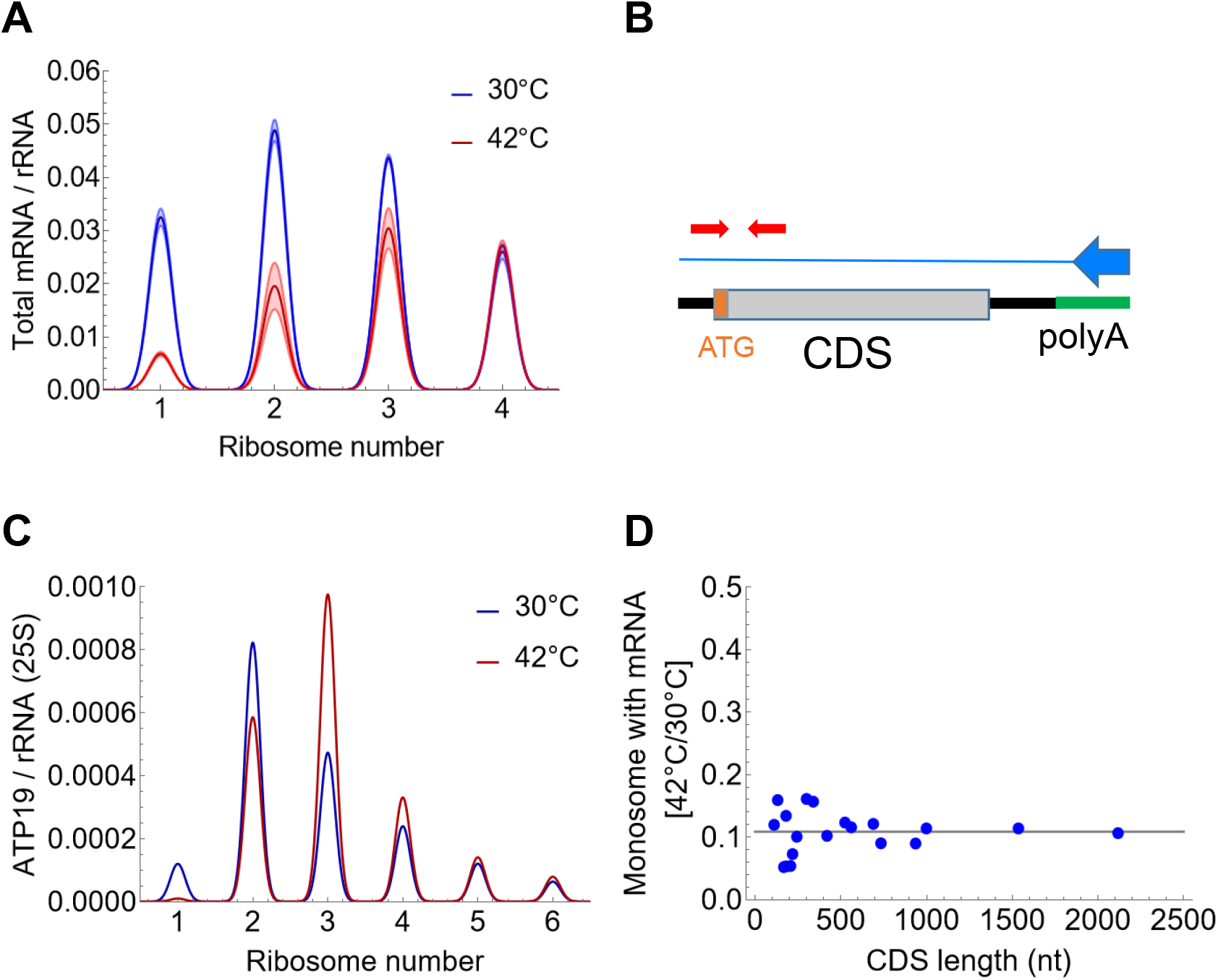
Estimation of the silent ribosomes using RNA-seq and RT-qPCR. **(A)** RNA-seq data obtained from RNA samples in which rRNA was not depleted. mRNAs and rRNAs (5.8S, 18S, and 25S) levels were summed in TPM units to calculate the ∑mRNA/∑rRNA ratio. The peak values of the dark and light bell-shaped curves denote the mean and standard deviation (SD) (*n* = 3 replicates). **(B)** Scheme showing primers used to measure full-length mRNAs. **(C)** The ATP19 mRNA/ 25S rRNA ratios measured by RT-qPCR. The peak N = 6 represents six or more ribosomes. **(D)** The fraction of monosomes containing full-length mRNAs at 42°C was estimated using RT-qPCR based on the following mRNAs: *OST4, PMP2, PMP3, ATP18, HOR7, ATP19, DAD4, SSS1, NHP6B, ECM19, MRPL50, ATP7, ATG12, PCL5, ATP4, ATP3, TDH3, ATP2* and *HSC82*.

This estimate likely represents an upper limit, as it does not consider the possibility that some monosomes are already devoid of mRNA at 30°C. This idea is supported by examining the relationship between ∑mRNA/∑rRNA ratios and ribosome numbers. Fractions with fewer ribosomes are expected to have higher ∑mRNA/∑rRNA ratios because larger ribosome assemblies, such as tetrasomes, associate more ribosomes with an mRNA compared to smaller assemblies like disomes. This trend was indeed observed at 30°C between disomes and tetrasomes. However, monosomes deviated from this pattern, as revealed by linear regression analysis (Fig. S1D). A comparison of the extrapolated value from the regression to the measured value indicates that 48% of monosomes are devoid of mRNA even at 30°C. Accounting for this adjustment reduces the estimate of mRNA-loaded monosomes at 42°C by half. The remaining ribosomes are considered silent.

We referred to ribosomes devoid of full-length mRNAs as silent ribosomes rather than vacant ribosomes, as our measurements cannot exclude the presence of short RNA fragments, which are undetectable by RNA-seq under the applied settings. Depending on whether extrapolation is applied to the monosome loading state at 30°C, silent ribosomes at 42°C are estimated to constitute 80% to 90% of the monosome fraction.

To validate these estimates, we used an alternative mRNA detection method. Specifically, we utilized RT-qPCR with oligo-dT-based reverse transcription and AUG-centered amplicons for the qPCR, ensuring the detection of poly(A)-tailed, full-length, translation-compatible mRNAs (Fig. 2B). mRNA levels were adjusted based on the measured PCR amplification efficiencies and subsequently normalized to 25S rRNA (*RDN25*).

We selected mRNAs with coding sequence (CDS) lengths ranging from 111 to 2118 nucleotides. For example, analysis of *ATP19* mRNA revealed a substantial reduction in the proportion of mRNA-loaded monosomes after heat shock (Fig. 2C). Silent ribosomes were estimated from the ratio of monosome propensities based on normalized mRNA levels at the two temperatures (see Methods, Estimation of the proportion of monosomes with full-length mRNAs under heat-shock using RT-qPCR).

At 42°C, 10.8 ± 3.3% (median ± standard deviation) of the monosomes were estimated to contain full-length mRNAs (Fig. 2D). The estimates from the specific mRNAs were distributed narrowly around the median and were independent of the CDS lengths (*r*_*S*_ = 0.007, P-value = 0.97). This finding—indicating ~90% silent ribosomes— aligns closely with the estimates based on RNA-seq samples that were not depleted of rRNA.

### Proteomic characterization of silent ribosomes

To characterize the composition of silent ribosomes, we analyzed the proteins in each fraction using mass spectrometry. Protein abundances were normalized to the average abundance of ribosomal proteins (RPS and RPL) (see Methods). The resulting distribution of normalized abundances is bimodal (Fig. S3A, B), with the smaller peak corresponding to the highly abundant ribosomal proteins, and the larger peak representing mostly the other less abundant associated proteins (Data S2). Heat shock causes only a minor shift in the distribution of proteins in tetrasomes but reduces the abundance of non-ribosomal proteins in monosomes, by approximately eight times (Fig. S3A, B). This substantial depletion of ribosome-associated proteins reinforces the translationally silent nature of these monosomes.

In order to characterize temperature-induced changes in the association of specific proteins, we defined two parameters: TICTRA (temperature-induced changes in total ribosomal association) and TICMA (temperature-induced changes in monosomal association). TICTRA reflects changes in protein association with all ribosomes, while TICMA specifically reflects those with monosomes, normalized against tetrasomal association (see Methods: Analysis of protein abundances in the ribosome profiling fractions).

Heat stress reduced the total ribosomal association of 21 proteins to less than half (TICTRA < 0.5), with translational initiation being the gene ontology term that was highly enriched among them (Pat1, Cdc33, Gcd11, Nip1, Prt1, Tif34, Tif4632, Xrn1; fold-enrichment = 35.4, false-discovery rate (FDR) = 1.14·10^-7^). Conversely, 37 proteins displayed increased ribosomal association (TICTRA > 2). These proteins typically participate in de novo post-translational protein folding (Ssa1, Ssa3, Ssa4, Sis1, Hsp104; fold-enrichment = 34.1, FDR = 2.95·10^-4^) and cellular response to heat. These findings suggest that while recruitment of translational initiation factors decreases under heat stress^16^, the translated proteins are increasingly chaperoned.

To analyze the monosomes specifically, we calculated TICMA. Most proteins exhibited TICMA < 1, confirming reduced monosomal association at 42°C (Fig. 3A). We analyzed the function of the proteins with TICMA < 0.1. This threshold helps identify proteins that exhibit a tenfold or greater depletion, corresponding to the extent of mRNA depletion relative to rRNA in silent ribosomes. Translation initiation factors and chaperones, despite their contrasting total ribosomal association patterns, were both significantly depleted in silent ribosomes (Table 1). Under normal conditions (30°C), the Ssa3 chaperone binds uniformly across the ribosome configurations, suggesting a stoichiometric association of this chaperone to the translating ribosomes (Figs. 3B, S4A). However, heat shock enhances the binding of Ssa3 only to the polysomes leading to a relative depletion of the monosome. Conversely, Cdc33 (eIF4E), a translation initiation factor binds around 12 times more to the monosomes under normal conditions (Figs. 3C, S4B). This indicates that upon the pioneer round of translation initiation, monosomes constitute the dominant ribosomal configuration^3^. Heat shock abolishes the preferential binding of Cdc33 to monosomes, leading to a relative reduction in its association with monosomes at 42°C. Although the total association of the two proteins changes in opposite directions, these shifts collectively lead to a relative depletion of both Ssa3 and Cdc33 in monosomes during heat shock.

**Table 1.** Enriched biological processes in proteins with TICMA < 0.1. GOs with at least 4 protein hits are shown with a fold enrichment > 9.

| Gene ontology<br>(Biological process) | Fold En-<br>richment | P-value | FDR | #P<br>GO | Genes |
| --- | --- | --- | --- | --- | --- |
| 3'-5' exonucleolytic<br>nonsense-mediated<br>decay** | 20.9 | $2.0 \cdot 10^{-5}$ | $2.09 \cdot 10^{-3}$ | 8 | SKI2, SKI3, SKI8, NAM7 |
| 'de novo' post-translational protein<br>folding | 13.9 | $4.7 \cdot 10^{-8}$ | $2.02 \cdot 10^{-5}$ | 24 | CCT3, CCT5, HSP104, SIS1,<br>SSA1*, SSA3*, SSA4",<br>SSD1* |
| tRNA<br>aminoacylation for<br>protein translation | 11.3 | $9.0 \cdot 10^{-9}$ | $1.06 \cdot 10^{-5}$ | 37 | ARC1 DED81, DPS1,<br>GLN4, GUS1, HTS1, KRS1,<br>MES1, VAS1, YHR020W |
| Glycolytic process | 10.4 | $1.6 \cdot 10^{-6}$ | $1.75 \cdot 10^{-3}$ | 24 | CDC19, ENO1, GPM1,<br>HXK1, TDH3, TPI1 |
| Cytopl. translational<br>initiation | 9.5 | $1.4 \cdot 10^{-4}$ | $7.9 \cdot 10^{-3}$ | 22 | NIP1, PAT1, PRT1, TIF3,<br>TIF34 |

\*These genes also constitute the “SRP-dependent co-translational protein targeting to membrane, translocation” (Fold enrichment = 12.9).
\*\*Nuclear-transcribed mRNA catabolic process, 3'-5' exonucleolytic nonsense-mediated decay

**Figure 3.**
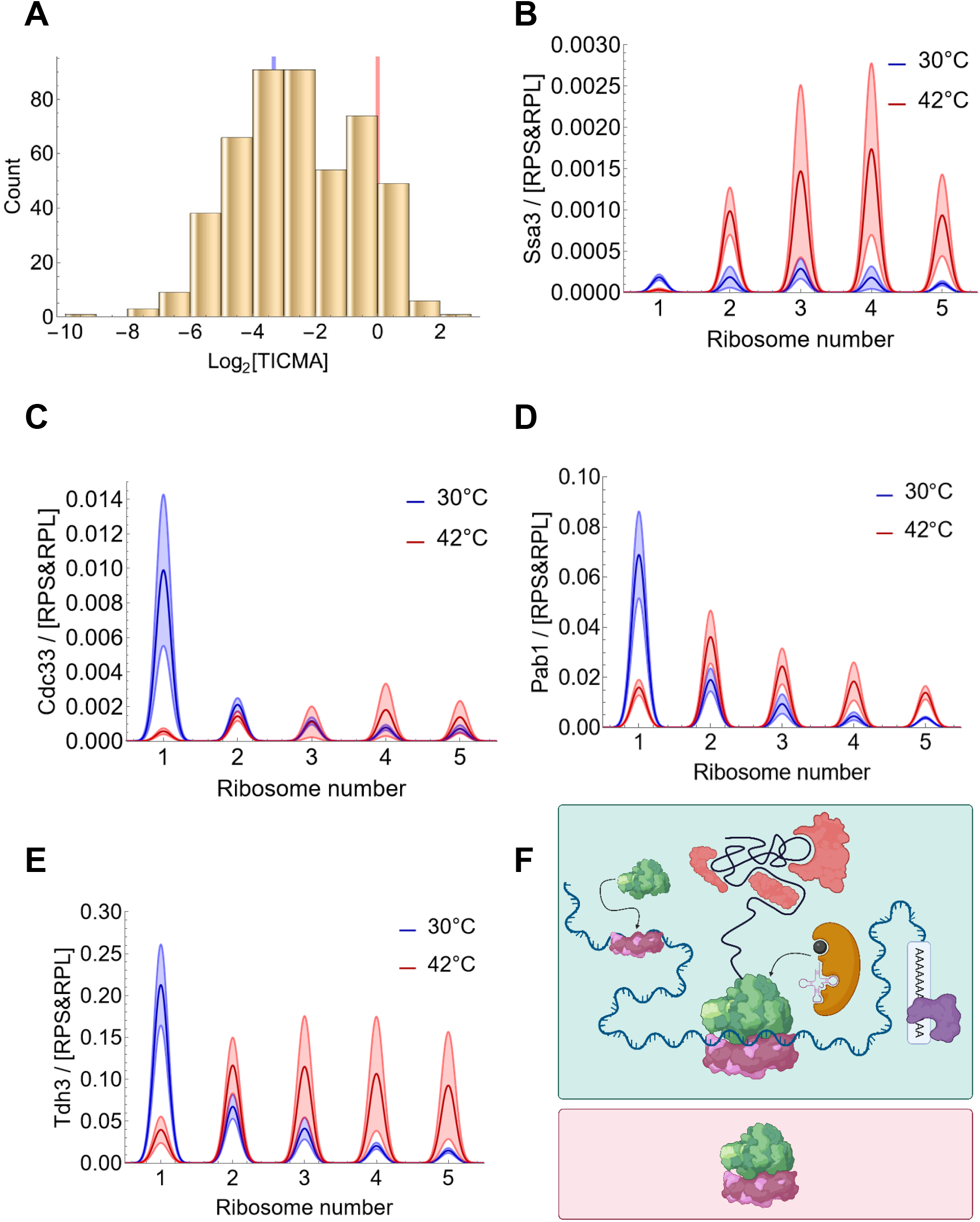
Proteomic analysis of the polysome profiling fractions. **(A)** Distribution of TICMA values of the proteome. Proteins with TICMA < 0.1 (blue line) were analyzed for enrichment. **(B-E)** The peak values of the dark and light bell-shaped curves denote the mean and standard deviation (*n* = 4 biological replicates) of the normalized protein abundances, respectively. Normalization was performed using the average of Rps and Rpl ribosomal proteins in each of the indicated fractions of the polysome profiling. The respective TICTRA and TICMA values of the proteins are indicated in parentheses: Ssa3 (5.438, 0.018), Cdc33 (0.432, 0.025), Pab1 (1.029, 0.056), and Tdh3 (1.319, 0.035). **(F)** Scheme of the monosome-associated proteins at 30°C (top panel) and at 42°C (bottom panel) based on the enrichment analysis.

A further biological process depleted in silent ribosomes is also related to translation: tRNA amino-acylation, a process that supplies amino acids for translation. The anchoring of the enzymes catalyzing the aminoacylation of the tRNAs to the ribosomes is mediated by rRNA segments^17,18^. This anchoring may enhance translation through channeling or other extra-translational processes^19^. Thus, the silent monosomes are depleted of proteins participating in several aspects of translation, such as translational initiation or co-translational folding, including targeting to ER, and aminoacylation of the tRNAs (Table 1, Fig. 3F).

Unexpectedly, glycolysis is enriched among proteins with TICMA < 0.1. Many glycolytic enzymes have dual roles, including RNA-binding functions, highlighting their moonlighting function^20,21^. To investigate how these enzymes interact with the ribosomal assemblies, we analyzed the monosome bias of two classes of proteins at standard conditions: mRNA-binding proteins and ribosome-associated proteins.

Many mRNA binding proteins, such as the Ski complex, Pab1 and Cdc33, exhibit preferential binding to specific regions or ends of the mRNA. The Ski complex is involved in the 3’-to-5’ degradation of mRNAs^22^, while Pab1 binds the polyA tail at the 3’-end of the mRNAs, and Cdc33, a translation initiation factor, binds to the 5’-cap. All of these proteins are depleted in the silent monosomes (TICMA < 0.1). Proteins that bind to mRNA ends, such as the 5’ cap or 3’ poly(A) tail, are expected to exhibit non-stoichiometric ribosomal association because their binding is independent of the number of ribosomes translating the mRNA. Consequently, there is an inverse relationship (bias) between the number of ribosomes in a fraction and the abundance of mRNA-end-binding proteins in these fractions. At 30°C, Pab1 exhibits a 15-fold monosome bias (monosome/tetrasome ratio >> 1 at 30°C), while Cdc33 shows a 12-fold bias (Figs. 3C, D and S4B, C), confirming an inverse relationship. In fact, these biases exceed the expected four-fold ratio (tetrasome; *n* = 4), likely due to the dissociation of initiation factors, like Cdc33, from polysomal mRNAs during the early stages of translation.

In contrast, uniform binding across monosomes and polysomes was observed for stoichiometric ribosome-binding proteins, such as the chaperone Ssa3 (monosome bias = 1.02, Fig. S4A). Similarly, ER translocon proteins, Sec61 and Sec62, showed no monosome bias, with values near one (0.75 and 0.74, respectively). This uniform association indicates that each ribosome remains linked to a translocon, irrespective of the number of ribosomes translating the mRNA.

Glycolytic enzymes can, in principle, display a monosome bias if they bind mRNA in a sequence-specific manner or, alternatively, display no bias by binding uniformly to each ribosome or non-specifically to the mRNA. A strong 10-fold monosome bias is observed for the Tdh3 enzyme (Fig. 3E), a pattern mirrored by other glycolytic enzymes (Table 1, Data S3). This suggests that glycolytic enzymes bind directly to specific regions of the mRNA, rather than associating indirectly through ribosomes.

Given that both glycolytic enzymes and RNA-binding proteins, such as Pab1 and the Ski complex, are depleted in heat-shocked monosomes, silent ribosomes not only lack proteins involved in translation but also those directly associated with the mRNA.

TICMA less than 0.1 highlights depletion patterns, while TICMA > 1 reveals enrichment of proteins in the silent ribosomes (Fig. 3A). We found few ribosome associated proteins that were enriched in the silent ribosomes, such as the Mbf1 and Stm1. Mbf1 prevents frame-shifting in stalled ribosomes^23^, and Stm1 is enriched in vacant ribosomes under starvation condition^24,25^. Another protein, Lso2 also has a strong preference for hibernating ribosomes^26^. We could not calculate its TICMA ratio because no Lso2 was detected in the tetrasome fraction at 30°C.

### Heat shock reduces codon-dependent differential mRNA stability

Some of the RNA-binding proteins, particularly exoribonucleases, play a critical role in mRNA turnover. Xrn1, a 5’-to-3’ exonuclease, along with the Ski proteins, which associate with the 3’-to-5’ exonucleases, are notably depleted from silent monosomes (Fig. 4A, TICMA(Xrn1) = 0.078). Moreover, the overall association of Xrn1 with the total ribosomes decreases to one-third at 42°C (TICTRA = 0.36). Given that Xrn1 is a key determinant of mRNA degradation rates and codon-dependent differential mRNA stability, we hypothesized that the reduced association of this protein with mRNAs may slow degradation during heat shock. mRNA degradation rates increase with temperature, but less rapidly than predicted by van’t Hoff’s law^9,15^, suggesting that the reduced Xrn1 association with ribosomes could contribute to this phenomenon. Therefore, we investigated whether codon-dependent differential mRNA stability is similarly affected.

**Figure 4.**
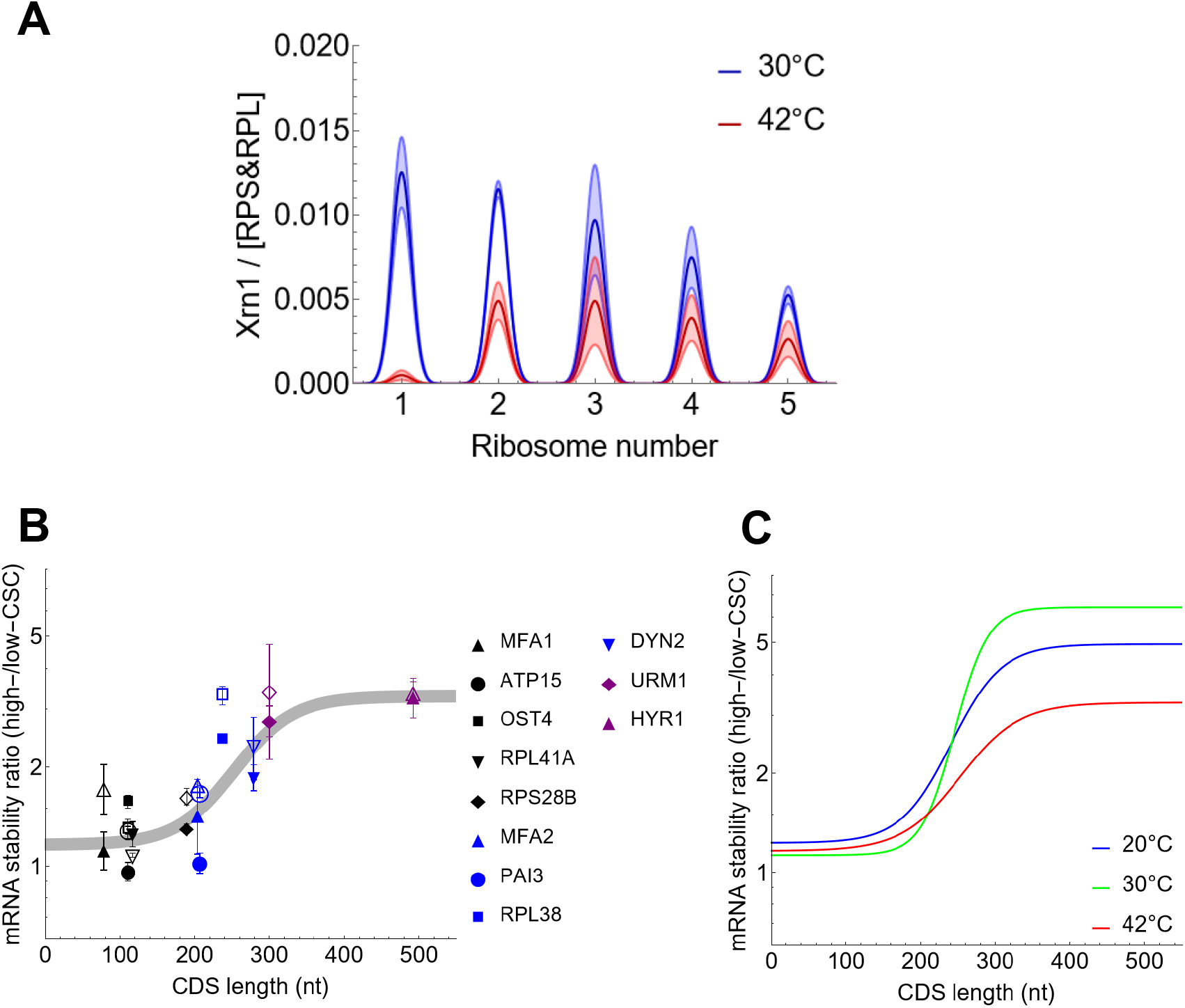
Differential stability of mRNAs upon heat-shock. **(A)** The peak values of the dark and light bell-shaped curves denote the mean and standard deviation (*n* = 4 biological replicates) of the normalized Xrn1 abundances, respectively. Normalization was performed using the average of Rps and Rpl ribosomal proteins in each of the indicated fractions of the polysome profiling. **(B)** Differential stability of recoded endogenous mRNAs. Differential stability is the ratio of the half-lives of the mRNA variants filled with high- and low-CSC codons. The filled symbols represent the mRNAs with original codon order, to which the equation was fit. The empty symbols correspond to the mRNAs swapped codon order, in which the positions of codon was pairwise swapped. **(C)** The differential mRNA stability curves were obtained by fitting equation 2 to the mRNA stability ratios measured at the indicated temperatures. The following parameters values ± standard error were fitted for 30°C: *b* = 1.12 ± 0.17, *DS*_*Max*_ = 5.28 ± 0.42, *L*_*C*_ = 263.9 ± 9.8 and *k* = 0.047 ± 0.010; experimental data from reference^11^; for 20°C *b* = 1.23 ± 0.42, *DS*_*Max*_ = 3.71 ± 0.89, *L*_*C*_ = 263.5 ± 25.1 and *k* = 0.030 ± 0.017; for 42°C: *b* = 1.16 ± 0.24, *DS*_*Max*_ = 2.12 ± 0.49, *L*_*C*_ = 272.9 ± 26 and *k* = 0.025 ± 0.015.

mRNAs encoding the same protein but with synonymous optimal or non-optimal codons exhibit markedly different half-lives, with optimal codons conferring longer half-lives and non-optimal codons leading to shorter ones^11^. The ratio of the half-lives of mRNAs with optimal and non-optimal codons is referred to as differential stability. Xrn1 plays a key role in differential stability, primarily through the rapid degradation of the mRNAs with non-optimal codons, but even stable mRNAs are stabilized slightly in *xrn1*Δ cells. The differential stability emerges above a threshold coding sequence (CDS) length^11^.

To investigate differential stability, we analyzed genes with CDS lengths both below and above this threshold, ranging from 100 to 500 nucleotides^11^. Codons in these mRNAs were systematically replaced with either optimal (high-CSC) or non-optimal (low-CSC) codons. Using an inducible gene expression system, we measured the half-lives of these mRNAs to assess the impact of codon usage and CDS length on mRNA stability. The half-lives were measured at 20°C and 42°C, and also compared to previously determined half-lives at 30°C^11^. A sigmoidal function was fitted to determine the maximal differential-stability and the CDS threshold length (Methods). The threshold length for the emergence of differential stability is remarkably stable across temperatures, consistently falling between 260 and 280 nucleotides (Fig. 4B). Potential feedback loops mediated by proteins expressed from the mRNAs do not change substantially differential stability as shown with mRNAs with swapped codon order (Fig. 4B, empty symbols).

Interestingly, while the threshold position was largely temperature-independent, the maximal differential stability was highest at 30°C, reaching 5.3, but was reduced by half at 42°C (2.1, Fig. 4C). This reduction is in agreement with the decreased ribosomal association of Xrn1 observed at 42°C. Additionally, the lower polysome propensity of mRNAs with longer CDSs at higher temperatures may contribute to this effect, as mRNAs exhibiting high differential stability tend to have high polysome propensity^11^.

### Estimation of mRNA turnover parameters

The results above indicate that heat shock not only reorganizes the ribosomal assemblies through the induction of silent ribosomes but also affects the final steps of mRNA turnover, as evidenced by changes in differential mRNA half-lives. To investigate this further, we aimed to estimate key parameters for the major steps of cytoplasmic mRNA turnover: the rate at which mRNAs associate with ribosomes and their degradation rates.

To achieve this, we developed a simple two-step mathematical model of cytoplasmic mRNA turnover (Supplementary Text). The steady-state solution of this model shows that rates of mRNA association with ribosomes can be calculated using two factors: the fractional occupancy of mRNAs (the proportion of mRNAs bound to ribosomes) and their half-lives. The fractional occupancy can be determined through polysome profiling using the previously obtained experimental data, while transcriptome-wide measurements of mRNA half-lives have to be determined to complete the analysis.

### Optimization of metabolic labelling across a broad temperature range

To measure the mRNA half-lives genome-wide, we performed metabolic labelling. To apply the method over a broad temperature ranges, we optimized the conditions for metabolic labeling. Measuring mRNA half-lives at elevated temperatures can be challenging, as the procedures required for different measurement methods may introduce additional stress^9,15,27^.

To label intracellular RNA, the metabolite 4-thiouracil, or its nucleoside derivative, 4-thiouridine, is typically applied at concentrations ranging from 0.5 to 5 mM^28^. To assess potential side effects of 4-thiouracil on mRNA half-life measurements, we exposed cells to 4-thiouracil and measured mRNA half-lives with the orthogonal genetic control assay at 42°C. Our results showed that while 5 mM 4-thiouracil does not affect the half-life of stable mRNAs, it significantly stabilizes short-lived mRNAs—by up to fourfold (Fig. 5A). This stabilization introduces a notable negative correlation between mRNA half-life and thiouracil-induced fold-change in half-lives. This effect can abolish much of the variations in mRNA stability since a 5fold difference in mRNAs stability (Fig. 5B, C) is reduced to approximately 1.25, a range comparable to measurement noise.

**Figure 5.**
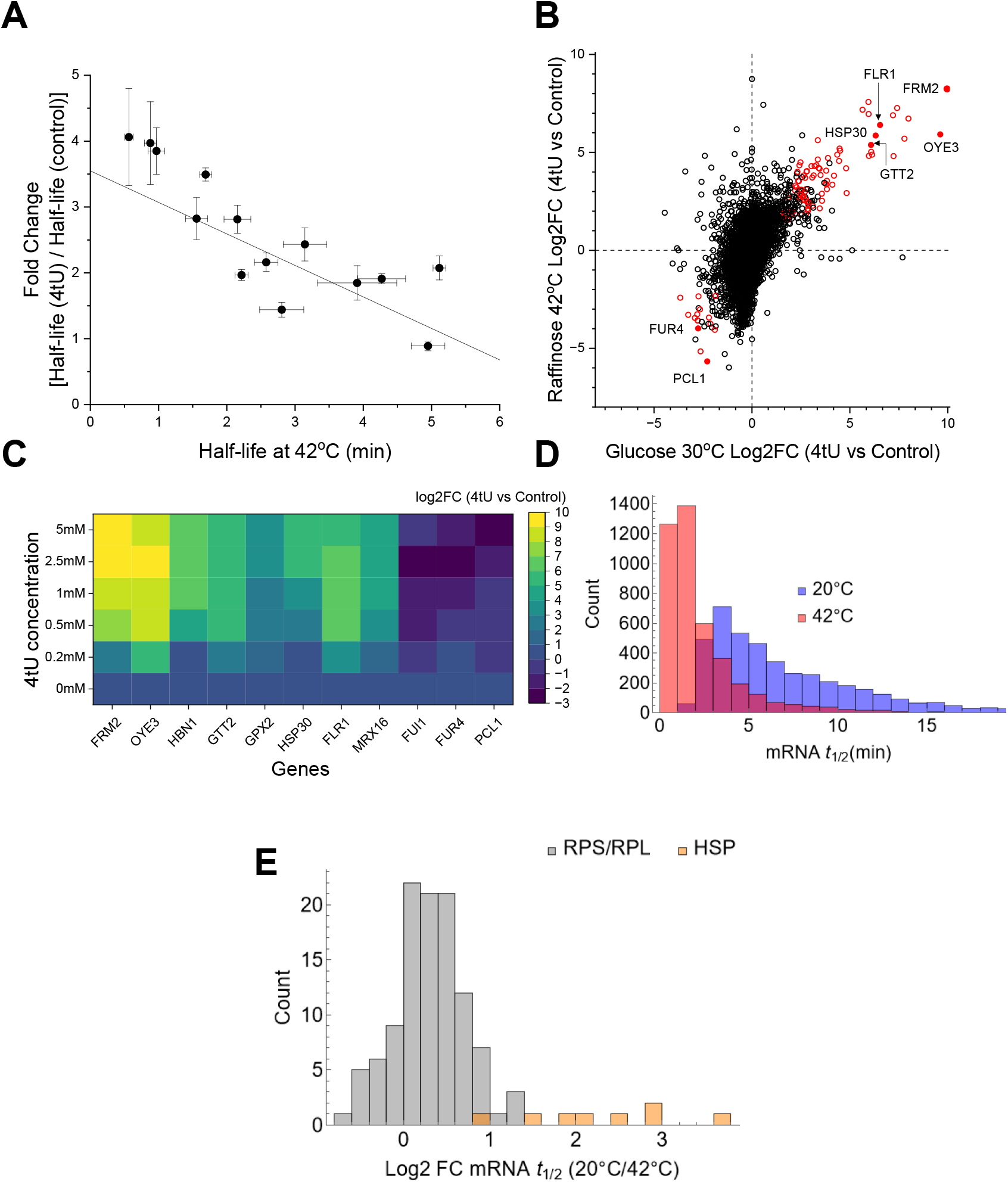
Metabolic labelling and measurement of mRNA half-lives. **(A)** Fold change in the half-lives of the representative set of mRNAs (Table S1) when heat shocked cells were treated with 5mM 4-thiouracil (4tU) for 6min as a function of mRNA half-lives (control). Error bars denote SEM (*n* = 3 replicates). **(B)** Log2 fold change (FC) in mRNA levels upon a 12 min treatment with 5mM 4tU at 30°C (glucose) and 42°C (raffinose). Significantly up- or downregulated genes (Adjusted P-value < 0.05) are represented with red circles. mRNAs used as markers for 4tU-treatment-related stress are represented by filled circles. **(C)** Fold change in the level of marker mRNAs indicative of 4tU-treatment-related stress at different concentrations of 4tU (30 min at 30°C, glucose). **(D)** Histogram showing the distribution of the half-lives measured with RNA metabolic labelling at 20°C and 42°C. **(E)** Histogram of temperature induced fold-changes (FC) in the half-lives of mRNA encoding ribosomal (*RPS* and *RPL*) and heat shock proteins (*HSP*).

To examine the temperature-dependent effect of 4-thiouracil, cells were exposed to 5 mM 4-thiouracil at 30°C and 42°C, followed by RNA-seq analysis. Most genes responded to 4-thiouracil similarly at both standard temperature and heat shock (Fig. 5B). The genes involved in the transport of uracil (*FUI1, FUR4*) were suppressed, whereas the highly induced genes were enriched for functions related to redox activity and cellular stress (Figs. 5B and S5; the nitroreductase-like oxidoreductases, *FRM1* and *HBN1*, the NADPH oxidoreductase, *OYE3*, the glutathione peroxidase *GPX2* and the glutathione S-transferase *GTT2;* Data S3).

To determine an acceptable range of 4-thiouracil concentrations, we analyzed the response of the above indicator genes across varying 4-thiouracil concentrations. Four of the eight highly induced genes exhibited maximal expression at 2.5 mM 4-thiouracil, while three others peaked already at concentrations exceeding 1 mM (Fig. 5C). To avoid triggering a full stress response, we opted to use a lower concentration of 0.5 mM for the subsequent metabolic labelling.

To assess the incorporation of the 4-thiouracil into the RNA, culture was sampled after metabolic labelling. The isolated RNAs was subject to biotinylation at the thiol moieties of the incorporated 4-thiouracil in the labelled RNA. In vitro transcribed 4-thiouracil labelled spike-in mRNAs were added to the samples to permit data normalization. Subsequently, the biotin conjugated labelled RNAs were separated from the unlabeled RNA by magnetic streptavidin beads. The bound mRNA was released by cleaving the disulfide bond with beta-mercaptoethanol. The eluted labelled mRNA was then precipitated and reconstituted for further analysis by qPCR and RNA-seq. RNA-seq was performed as described in the Methods. RNA levels (with TPM units) were normalized to spike-ins as in vitro controls (Fig. S6). Afterward, we divided the normalized values by the intron counts of selected mRNAs to act as in vivo control, a process we termed relative normalization (Fig. S7, Methods)^29^. From these values, we used the approach to equilibrium formula to estimate the half-lives. Since median based normalization increases the robustness of half-life estimation^11^, we scaled the half-lives of each replicate linearly to the half-lives of selected reference mRNAs measured by an independent metabolic labelling experiment quantified by qPCR. This absolute normalization yields the final half-life estimates (Data S4).

The resulting half-life distributions were skewed toward long half-lives (Fig. 5D). The correlation between the half-lives measured at 20°C and 42°C was high (*r*_S_ = 0.84), confirming that precise half-lives can be reliably obtained using metabolic labelling, even at high temperatures. The high correlation also suggests that the degradation program is relatively robust over a broad range of temperatures. Heat-shock protein mRNAs show a median half-life reduction of 5.2 when the temperature increases from 20°C to 42°C, compared to a reduction of 1.2 for ribosomal protein mRNAs (Fig. 5E). This is intriguing, as it indicates that mRNAs encoding heat-shock proteins do not gain stability during heat shock.

### Heat Shock Induces Distinct Translational Profiles in Ribosomal and Heat Shock Protein mRNAs

The median mRNA occupancy declines from 0.86 (at 30°C) to 0.77 (at 42°C), with heat-shock-repressed mRNAs—such as those encoding ribosomal proteins (RPS/RPL)—showing a more substantial reduction from 0.93 to 0.70 (Fig. 6A). Furthermore, the RPS and RPL mRNAs generally exhibit lower polysome propensities at 42°C (median PP = 4.1). In contrast, mRNAs encoding heat shock proteins typically show a higher polysome propensity (median PP = 11.3) (Fig. 6B). As a result, a weak positive correlation is observed between the induction ratio and the polysome propensity of mRNAs (Spearman correlation coefficient, *r*_S_ = 0.16, P-value = 2·10^-33^).

**Figure 6.**
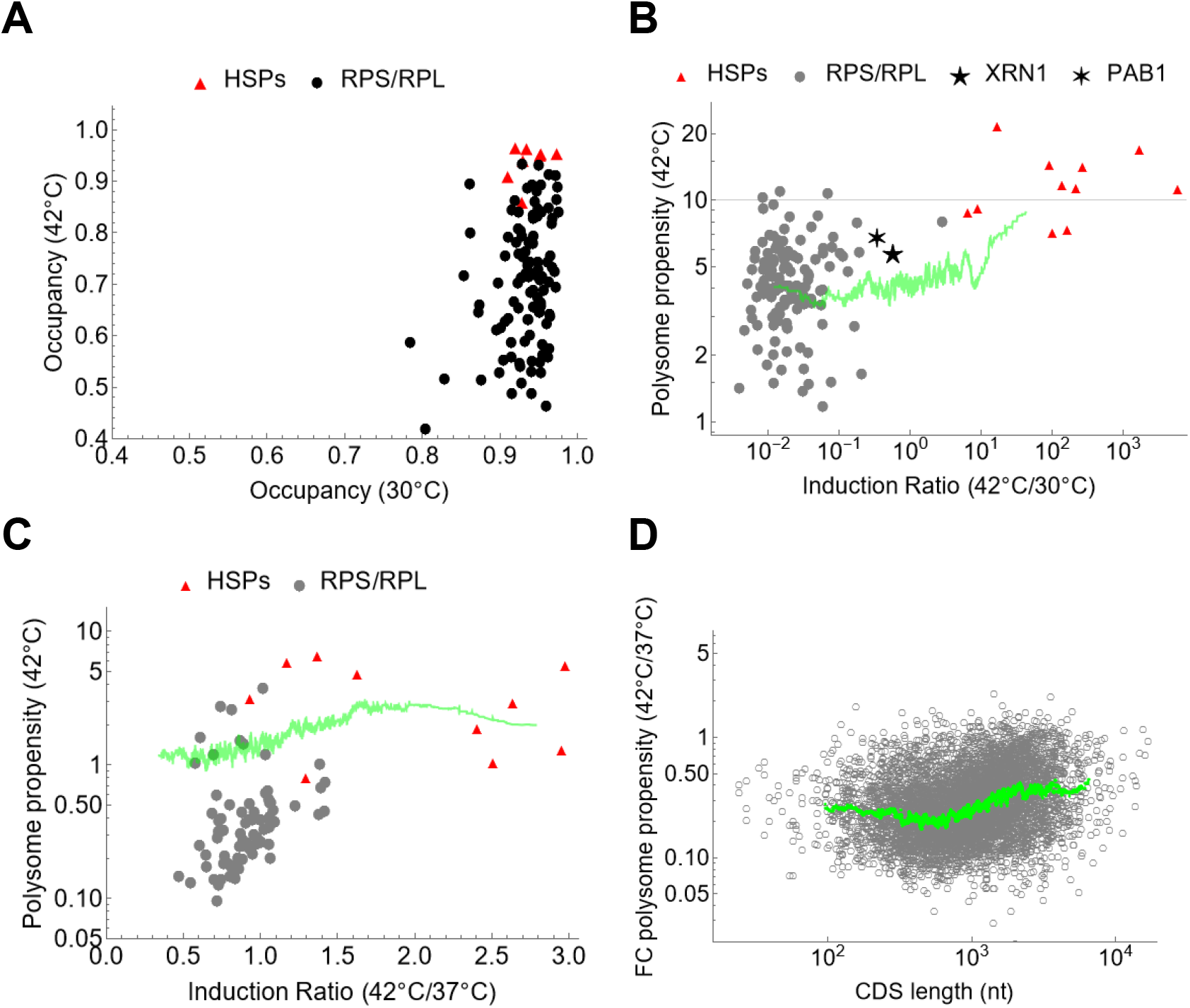
Temperature induced changes in the polysome propensity across the transcriptome. **(A)** Occupancy of mRNAs encoding heat shock (HSP) and ribosomal (RPS/RPL) proteins at 30°C and 42°C in yeast cells. **(B)** Polysome propensity of the mRNAs as the function of the induction ratio in yeast cells. The green line represents the moving average of 100 mRNAs. Individual data are shown for mRNAs encoding HSPs, ribosomal proteins (RPS/RPL) and XRN1, PAB1. **(C)** Polysome propensity of the mRNAs as the function of the induction ratio in murine macrophage cell line J774A.1. Further details as in (A). **(D)** The ratio of polysome Propensities as a function of CDS length in J774A.1 cells. The green line represents the moving average.

To evaluate the conservation of this response in mammalian cells, we analyzed the murine macrophage cell line J774A.1. Similar to yeast cells, mRNAs encoding heat shock proteins exhibited higher polysome propensities compared to those encoding ribosomal proteins (median PP = 3.0 and 0.37, respectively; Fig. 6C). Additionally, the overall reduction in polysome propensity in the transcriptome was more pronounced (median fold change = 0.26), (Fig. 6D, Data S5), in comparison to yeast (median fold change = 0.72). In macrophages, mRNAs with short coding sequences also exhibited a marked reduction in polysome propensity (Fig. 6D).

### Balance between the heat-shock induced changes in association and degradation rates

Next, we used the mRNA degradation rates to calculate the rates of mRNA association with the ribosomes based on the ribosome occupancy data. While the median degradation rate increases when the temperature is raised from 30°C to 42°C, the overall association rate decreases (Fig. 7A). A likely driver of this decline is the abundance of silent ribosomes.

**Figure 7.**
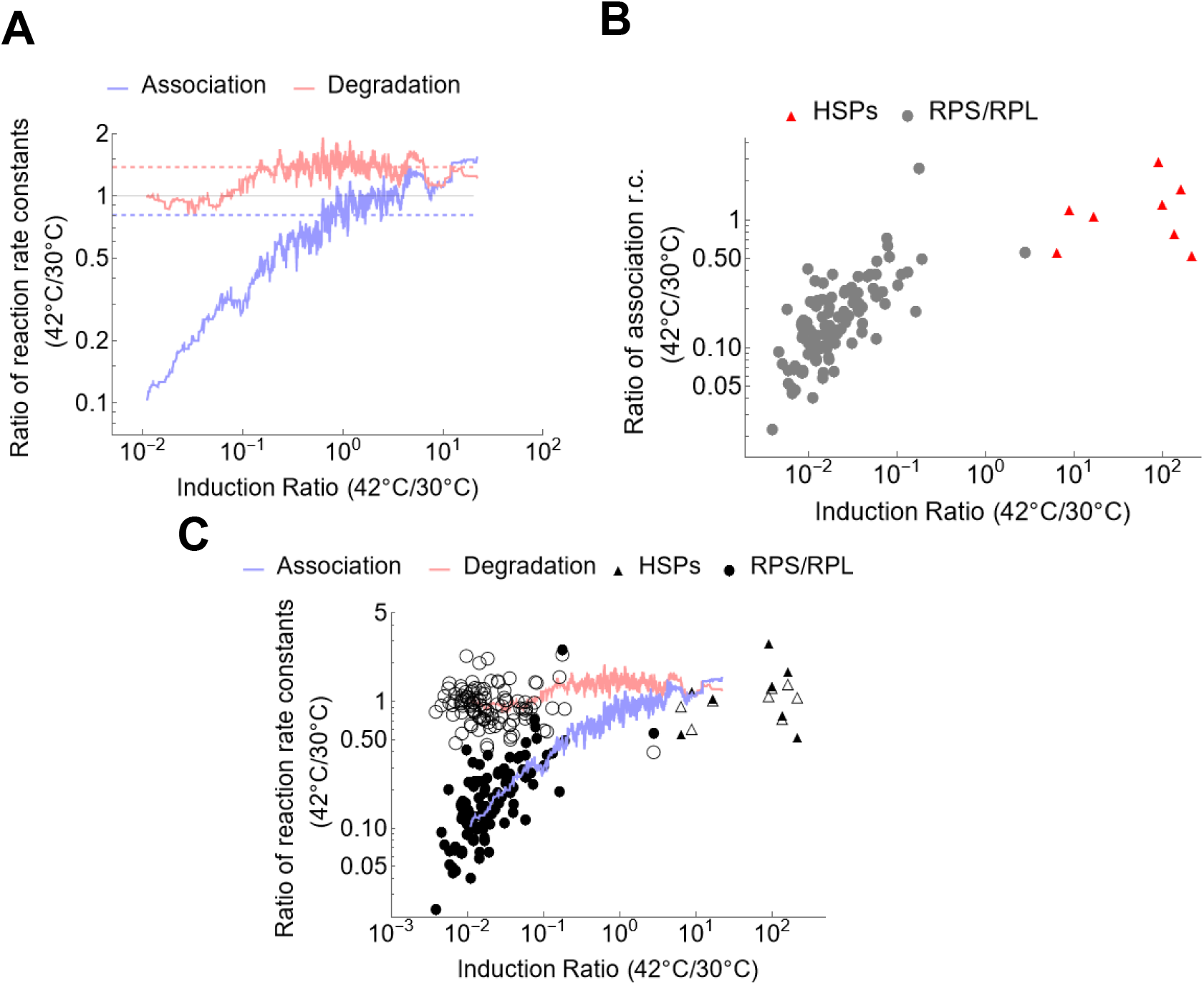
Heat-shock induced changes in association and degradation rate constants. **(A)** Solid lines represent the moving averages of the reaction rate constant ratios, while dashed lines indicate the transcriptome-wide medians of these ratios (median for the association = 0.80, and for the degradation = 1.37). (**B**) Ratios of the association rate constants for mRNAs encoding heat shock proteins (genes containing “HSP” in their names) and ribosomal proteins (genes containing “RPS” or “RPL” in their names). **(C)** Filled and empty symbols represent the ratios of association and degradation rate constants, respectively. Other details follow the conventions described in (A, B).

To examine the gene specific changes in association, we examined genes most suppressed or induced by heat shock, encoding ribosomal and heat-shock proteins, respectively (Fig. 7B). The association rate constant correlates positively with the induction ratio (Fig. 7A): mRNAs encoding suppressed ribosomal proteins show significant reductions in association rates, whereas those encoding heat-shock proteins maintain or even accelerate their association (Fig. 7B). Notably, no such correlation exists between degradation rates and induction ratios (Fig. 7A), indicating that heat shock exerts a more extensive regulatory influence on association rates than on degradation rates.

Interestingly, for highly induced heat-shock protein mRNAs, association and degradation rates are approximately balanced (Fig. 7C). In contrast, for ribosomal protein mRNAs, association rates lag behind degradation rates.

This imbalance allows heat-shock proteins to sustain steady expression and function while ribosomal protein expression is set to decline, preventing cells from sustaining growth under stress.

## DISCUSSION

Our results confirm that heat stress causes a substantial increase in the monosome peak in the polysome profile, a phenomenon consistently observed across eukaryotic organisms^12,30,31^. The accumulation of monosomes in heat shock has been typically interpreted as a result in the blockage of translational initiation or elongation^12,31^. However, the increase of monosome fraction after the temperature shift in yeast cells is not accompanied by a corresponding accumulation of monosomal mRNAs. We attributed this discrepancy to the presence of ribosomes devoid of mRNAs, which we refer to as silent ribosomes.

Silent ribosomes are commonly observed during nutrient stress conditions, such as starvation, as well as in processes like sporulation that occur in response to such stress^24^. These ribosomes, often referred to as inactive, dormant, or vacant, form when translation terminates. Normally, after translation termination, ribosomal subunits dissociate and participate in subsequent rounds of protein synthesis. However, under translation-inhibiting stress conditions, such as starvation, free ribosomal subunits can re-associate to form a large pool of non-translating 80S ribosomes. In yeast, these ribosomes are stabilized by the Stm1 “clamping” factor, while in mammalian cells, they are stabilized by SERBP^25^. The dissociation of these inactive ribosomes, which is crucial for the resumption of translation, is facilitated by specific proteins when yeast cells recover from starvation stress^32^.

Our results indicate silent ribosomes can form in yeast cells even in the absence of starvation, when the cells are exposed to elevated temperatures. Estimates based on RNA-sequencing of non-depleted samples and RT-qPCR of full-length mRNAs indicate that 80-90% of the ribosomes in the monosome fractions are devoid of mRNAs. Proteomic analysis reveals that these silent ribosomes are depleted of proteins involved in post-translational protein folding and key translational processes, such as translation initiation and tRNA amino-acylation. Additionally, they are depleted of RNA-binding proteins, including the poly(A)-binding protein Pab1 and certain glycolytic moonlighting proteins involved in RNA-binding. Interestingly, the composition of silent ribosomes represent the complement of stress granules. While stress granules are enriched with initiation factors, Pab1, and ribosome-free mRNA, silent ribosomes are depleted of these components. Stress granules, composed of mRNAs stalled in the process of translation initiation, typically form at higher temperatures, such as 46°C^33,34^. Pre-stress granules, containing most but not all components of fully formed stress granules, can assemble already at 42°C^33,35^. As a result of the residual translation, polysomal mRNAs persist during heat shock; however, actively translating monosomes are largely absent, having been partitioned into silent ribosomes lacking full-length mRNAs and pre-stress granules containing mRNA.

Although ribosome-associated proteins are largely depleted in silent ribosomes, some are enriched. Stm1, a clamping protein, stays bound to monosomes during heat shock, while proteins such as Mbf1 and Lso2 are present in higher amounts. Lso2, in particular, is a conserved ribosome-bound protein crucial for yeast translational recovery^24,36^. Thus, our study reveals that silent or hibernating ribosomes are not confined to starvation conditions but are an integral part of the heat-shock response, as well.

Monosomes are particularly abundant in neurons, often translating mRNAs encoding proteins in synaptic membranes^4^. Such proteins play an important role in cell fate specification^37–39^.

Interestingly, the Xrn1 exonuclease associates less with mRNAs under heat shock conditions. As Xrn1 is a key effector of RNA decay, this reduced association is expected to diminish codon-optimality-dependent differences in mRNA half-lives^35^. The narrowing of codon-optimality-dependent differential half-lives may also partly result from the accumulation of silent ribosomes since their accumulation leads to an increase in free mRNAs, which exhibit codon-independent neutral half-lives.

Our findings suggest that silent ribosomes are an important link in the regulatory mechanisms underlying the heat-shock response. The induction of silent ribosomes correlates with a lower-than-expected increase in mRNA association rates with ribosomes during heat shock, potentially due to a lack of sufficient ribosomes available for translation. Notably, mRNAs encoding ribosomal proteins exhibit a dramatically reduced association rate, resulting in decreased production of ribosomal proteins. Few mRNAs, primarily those encoding heat shock proteins, display balanced changes in ribosomal association and degradation rates. This balance allows for steady-state expression of heat shock proteins. In contrast, the decay rate of mRNAs encoding ribosomal proteins surpasses their ribosomal association rates, which will leads to a decline in ribosome biogenesis and translation.

Interestingly, while mRNAs encoding heat shock proteins are transcribed more rapidly and associate more quickly with ribosomes during heat shock, their half-lives do not increase. This can be attributed to the dynamic nature of the heat-shock response, as cells must adapt swiftly to temperature fluctuations. Rapid mRNA turnover ensures a prompt response to fluctuations even at elevated temperatures. Indeed, gene expression during heat shock exhibits a pulse-like pattern, possibly mediated by a feedback loop^40,41^. This transient response underscores the importance of maintaining rapid mRNA turnover to support cellular adaptation at high temperatures. The increased transcription and association rates, on the other hand, ensure the robustness and intensity of the heat shock response.

## RESOURCE AVAILABILITY

### Lead contact

Requests for further information and for resources and reagents should be directed to and will be fulfilled by the lead contact, Attila Becskei.

### Materials availability

This study did not generate new unique reagents.

### Data and code availability

RNA-seq data is deposited at the GEO database of the NCBI for polysome profiling of yeast samples at 42°C with rRNA depletion (GSE289484). The polysome profiling of yeast samples with no rRNA depletion (DOI: 10.5281/zenodo.14860597), and polysome profiling of macrophage with rRNA depletion, at 37°C (DOI: 10.5281/zenodo.14860705) and at 42°C (DOI: 10.5281/zenodo.14864741) are deposited at ZENODO. MATLAB was used for the calculation of half-lives from the metabolic labelling experiments, which is available from the lead contact upon request.

## ACKNOWLEDGEMENTS

We thank Samuel Mondal and Simone Faravelli for the help with the processing of RNA-seq data, Sandrine Wallerich for experimental help, Phillippe Demougin (Genomics Facility, University of Basel / BSSE) for the preparation of Illumina libraries, and Gowtham Thambra for helpful discussions. J774A.1 murine macrophage cells were kindly provided by Jean Pieters. This work was supported in part by the Swiss National Science Foundation (310030_185001).

## AUTHOR CONTRIBUTIONS

S.R. and A. B. designed the experiments. S.R., N.S., N.M. and K.E.F. performed the experiments. A. B., S. R. and M.Z. wrote the manuscript. A.B. and S.R. analyzed the data.

## DECLARATION OF INTERESTS

The authors declare no competing interests.

## DECLARATION OF GENERATIVE AI AND AI-ASSISTED TECHNOLOGIES

During the preparation of this work, the authors used ChatGPT for text corrections. After using this tool, the authors reviewed and edited the content as needed.

## SUPPLEMENTAL INFORMATION

Supplementary Text, Table S1, Figures S1-S7

Data S1. Polysome propensity and occupancy of yeast mRNAs at 42°C.

Data S2. Protein abundances in the Polysome profiling fractions.

Data S3. Differential expression of yeast mRNAs due to 4-thiouracil perturbation.

Data S4. Yeast mRNA half-lives estimated with metabolic labelling at 20°C and 42°C.

Data S5. Polysome propensity and occupancy of macrophage mRNAs at 37°C and 42°C.

## METHODS

### Transcriptomic analysis of the effect of 4tU at different temperatures in yeast cells

Yeast cells (BY4742α) were grown overnight in complete synthetic medium (Complete Supplement: Formedium #DCS0019, Yeast Nitrogen Base: Formedium #CYN0410) in 2% raffinose and 0.005% glucose for the 42°C samples or 2% glucose for the 30°C samples) at 30°C. The overnight cultures were refreshed to an optical density at 600nm (OD600) of 0.15 and grown at 30°C until they reached the mid-log phase, with final OD600 of 0.5, for further experiments. For the subsequent standard-temperature experiment, cells were treated with 5mM 4tU (or vehicular control DMSO) for additional 12 min and collected. For the heat-shock experiment, the cells were filtered out onto a 0.22µm membrane and transferred to a 42°C medium. 18min after the temperature shift, the cells were treated with 5mM 4tU (or DMSO). 12 min later, the cells were collected. Cells were collected in chilled 50% methanol. The samples were spun down for 10min at 2500g. Lysis was carried out using MPBio Lysing Matrix Y tubes in a FastPrep-24™ 5G bead beating grinder and lysis system. RNA was extracted with RNeasy Mini Kit (Qiagen # 74104). Two biological replicates were used for RNASeq. rRNA was depleted with Illumina RiboMin Gold. Libraries were generated using Illumina TruSeq Stranded Total RNA. Barcoded libraries were sequenced at single ends in Novaseq6000 (Illumina) flow cell type S1. The reference transcriptome S288C (assembly R64) was used for mapping (Data processing of RNA-seq data). Differential expression analysis of genes was carried out with DeSeq2^42^.

### Polysome profiling of yeast cells and transcript quantification

For cell culturing, *S. cerevisiae* BY strains were grown at 30°C in complete synthetic media (Complete Supplement: Formedium #DCS0019, Yeast Nitrogen Base: Formedium #CYN0410) (with 2% raffinose and 0.005% glucose) until the mid-log phase (OD600 = 0.6) at 30°C. For the heat-shock experiment, cells were transferred to a 42°C medium upon filtration with a 0.45µm filter and resuspension. They were grown for an additional 30 min before sampling.

For the sampling, cells were collected by addition of 1 volume of culture to 2 volumes of chilled medium (0°C) containing cycloheximide (CHX) (ThermoScientific #357420050) at 1.5x (150 µg/mL). The resulting mixture was immediately submerged in a −20°C bath (1:1 mix of ethanol and water) for two minutes (so that the final temperature is less than 10°C) The cells were spun down at 4000rpm (3000g) for 10min in a centrifuge at 4°C. The supernatant was removed and the pellet was immediately subjected to lysis. This cycloheximide pre-treatment combined with cooling was carried out to prevent ribosome runoff^43^ as well as the cycloheximide-induced transcriptional upregulation of genes involved in translation^44^. Since these upregulated transcripts are not translated, a lower than expected fractional occupancy would be observed for these mRNAs.

The cell lysis and gradient preparation was performed as previously described^11^.

For the ultracentrifugation, the A_254_ of the lysate was measured by mixing sample (or blank with Polysome Lysis Buffer) with water in a 1:20 volumetric ratio. 2-4 A_254_ unit equivalents of the sample was loaded onto the 10-50% sucrose gradient. Ultracentrifugation was carried out with a SW41Ti rotor (swing bucket) at 40000rpm (equivalent to 275,000g) for 2h at 4°C. The polysome profile was generated with BioComp™ Polysome Profiling machine. 25 fractions (450µL each) corresponding to different positions in the centrifuge tube were collected.

For RNA purification, the fractions were pooled according to the peaks of A_254_ polysome profile (free fraction, 40S, 60S, monosome, etc.), and in-vitro transcribed spike-in RNAs were added to each pool to serve as a volumetric control. Subsequently, the pools were subjected to phenol-chloroform extraction (Phenol: chloroform: isoamyl-alcohol = 25:24:1). Precipitation was carried out with 2.5 volumes of ethanol at −80°C overnight with 2µL of GlycoBlue (ThermoScientific #AM9516) as co-precipitant. RNA was reconstituted with 20µL of nuclease free water (ThermoScientific #AM9932) containing RNasIN (Promega #N2611) at 1µL/mL (40U/mL). This RNA was further purified using RNeasy MinElute Cleanup Kit (Qiagen #74204) with on-column DNase digestion. The purified RNA was then used for qPCR or RNA-Seq.

For qPCR based quantification, reverse transcription was carried out with SuperScript IV (ThermoScientific #18090200), using random hexamers (Promega #C1181) as per protocol and qPCR was then carried out (Roche LightCycler ® 480, KAPA SYBR Fast #KK4611). mRNA readouts were normalized to the geometric mean of the spike-ins.

For RNA-Seq, the ribosomal RNA was depleted with QIAseq FastSelect–rRNA (Qiagen). This depletion step was not applied when rRNA was quantified (non-depleted samples). cDNA libraries were prepared with Stranded SMART-Seq (Takara) by random priming. 10 and 13 cycles of amplification were performed in the first and second rounds, respectively. Size-selective purification of DNA fragments was performed with AMPure Beads as specified in Takara. The barcoded libraries were sequenced in Novaseq6000 (Illumina) flow cell type S1 resulting in single-end reads of 100-bp length.

### Polysome profiling of mammalian cells and transcript quantification

For cell culturing, J774A.1 murine macrophage cells were grown at 37°C in DMEM high glucose with 10% FBS and L-glutamine until ~80% confluence. For the heat shock experiment, the cells were transferred to medium at 42°C. They were grown for an additional 1h before sampling.

For the sampling, cells were collected by scraping in PBS supplemented with cycloheximide (CHX) (ThermoScientific #357420050) at 200µg/mL. Then the mixture was spun down at 500g for 3min in a centrifuge at 4°C. The supernatant was removed and the pellet was immediately subjected to lysis.

For the lysis, cells were dissolved in 1X mammalian Polysome Lysis Buffer [20mM Tris-HCl (ThermoScientific #AM9856), 10mM MgCl_2_ (ThermoScientific #AM9530G), 1mM DTT (ThermoScientific #707265ML), 50mM KCl (ThermoScientific #AM AM9640G), 200µg/mL cycloheximide (ThermoScientific #J66004-XF), 2 cOmplete EDTA free Protease Inhibitor (Sigma #COEDTAF-RO)/100mL, 16U/mL RNasIN (Promega #N2611), 1% v/v Triton X-100 (Fluka #93418)]. Lysis was carried out by passing the dissolved cells through a 23G syringe 10 times, followed by 10 cycles of bead-based lysis using Zirconia/Silica beads (Biospec #11079105z). The lysate was then clarified by a centrifugation for 5min at 10000g.

For the gradient preparation, 10% and 50% w/v sucrose (ThermoScientific #036508-30) solutions were prepared in 1x mammalian Polysome Gradient Buffer [as mammalian Polysome Lysis Buffer but without Triton X-100]. The 10% sucrose was loaded on the centrifuge tubes (Beckman C14293/344059 Ultraclear 13.2mL [compatible with SW41Ti rotor]). Subsequently, the 50% solution was loaded at the bottom of the centrifuge tube with a 20mL syringe having a long cannula attached to its mouth. The tubes were then covered with caps (BioComp #105-414-1). The gradient was formed by high angle rotation, using the 10%-50% short cap (rate zonal) protocol in the BioComp™ Gradient Station machine.

The ultracentrifugation, the RNA purification and the qPCR based quantification were performed as with yeast.

For RNA-Seq, cDNA libraries were prepared with Illumina TruSeq Stranded kit with Ribozero Gold rRNA depletion. Barcoded libraries were sequenced in Novaseq6000 (Illumina) flow cell type S4, PE101, resulting in paired-end reads of 100-bp length.

### Processing and analysis of RNA-seq data

For the data processing, single-end and paired-end sequence reads were trimmed for adaptor sequences using TrimGalore (v0.6.5) with --illumina parameter. Trimmed sequence reads were mapped, and transcript per million (TPM) values were obtained with Salmon (v1.9.0) ^45^. Specifically, Salmon’s index was created from National Center for Biotechnology Information (NCBI)’s reference transcriptome of yeast (S288C, assembly R64) and mouse (C57BL/6J, assembly GRCm39). TPM quantification was performed with salmon quant.

For the data analysis of the polysome profiling, the transcript TPM values were divided by the spike-in TPM values to obtain normalized transcript abundances. Subsequently, the normalized abundances were multiplied by the pooling factors. The pooling factor is equal to the number of fractions supplemented with spike-ins that were combined into a pool and sequenced. The pools typically represent a specific mRNA-ribosome configuration (e.g. monosome or tetrasome, etc.). The pooling factor is one if the spike-ins were added after the pooling.

Genes with zero counts in more than seven sequenced pools were removed; these were mostly noncoding RNAs, weakly expressed genes or short mRNAs. Furthermore, genes were filtered out if the polysome propensity had a coefficient of variation (CV) larger than 0.8.

Relative total mRNA values were calculated by dividing the total amount (sum of all fractions) of each mRNA by the median of the total values in the transcriptome. Relative total mRNA values were used to calculate the induction ratio of each mRNA measured at two different temperatures.

### Estimation of the proportion of monosomes with full-length mRNAs under heat-shock using RT-qPCR

RNA was extracted from Polysome profiling fractions as described earlier. Reverse transcription was carried out with SuperScript IV (Thermo #18090050) using oligodT_15_ (Promega #C1101) at a final concentration of 25µg/mL (5µM) along with specific RT primers for the spike-ins. The reverse transcription step was carried out at 50°C for 1h. From the resulting cDNA, specific transcripts were probed using qPCR with each forward primer encompassing the start codon of the particular transcript. This enabled us to assess the intensities of select transcripts that contained both the polyA tail and the start codon, i.e they are full-length. Each mRNA was normalized the spike-ins, yielding mRNA_i_. In order to estimate ratio of mRNA to the ribosome, the mRNA(rR)_*i,N*_ = mRNA_i_ / 25S rRNA ratios were calculated for each mRNA *i*, in each ribosome configuration (*N*).

25S rRNA abundances were obtained in a separate RT-qPCR experiment with gene specific primers corresponding to the reverse primer of the primer pair was used for the qPCR. Normalization was carried out by the spike-ins to get the intensities of the 25S rRNA. 25S (present in the large ribosomal subunit) rather than the 18S rRNA (present in the small subunit) was used in order to prevent signal from the scanning 40S ribosomal subunits that could be associated with mRNAs in each peak of the polysome profile.

The estimate for the monosomes with full-length mRNAs at 42°C, the ratio of monosome propensities at 42°C and 30°C was calculated for each mRNA_i_:

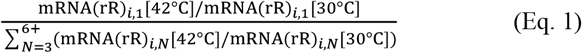

The monosome propensity is the inverse of the polysome propensity.

### PAGE analysis of rRNAs

Yeast and macrophage cells were lysed and polysome profiling was performed. ⅒ of the lysate used for the polysome profiling was preserved as a sample for total RNA. Fractions were collected and RNA was extracted as described in the polysome profiling methods. Equal volumes of RNA, along with Low Range ssRNA ladder (NEB #N0364S) as marker, were heat denatured by incubation at 90°C for 10min followed by snap-chilling. The samples were then loaded onto an 8% PAGE 8M Urea Gel. The RNA was visualized by incubating the gel for 10 min with SYBR Gold (Invitrogen # S11494) in TAE buffer (ratio of 1:10000). Band intensities were measured using FIJI, with background subtraction performed by measuring the intensity of gel regions devoid of visible bands.

### Determination of codon-dependent differential half-lives

mRNA half-lives were determined using the multiplexed gene control method as previously described^11^, with adjustments for different temperatures. In this method, strains containing gene expression cassettes encode the mRNAs of interest under a doxycycline-controllable promoter. Upon doxycycline addition, gene expression is repressed, and the ensuing decay profiles are analyzed with nonlinear regression to determine the mRNA half-life.

Briefly, *Saccharomyces cerevisiae* strains isogenic to BY4743 were grown in synthetic complete medium containing 2% raffinose and 0.005% glucose.

For the 20°C growth condition, cultures were initially grown at 30°C for 6–8 hours, followed by a temperature reduction to 20°C and overnight growth. Cultures containing the individual strains were then pooled to an OD600 = 0.35 and grown at 20°C until reaching an OD600 ~ 0.5, when doxycycline was added. For the heat-shock condition, overnight cultures were grown at 30°C. Cultures with the different were then pooled to an OD600 = 0.15 and grown at 30°C until reaching an OD600 ~ 0.5. Cells were filtered through a 0.45-µm membrane, and the filter was transferred to a fresh Erlenmeyer flask containing preheated medium (42°C). Cultures were incubated at 42°C for 30 minutes, after which doxycycline was added.

Transcriptional inhibition of tTA-controlled mRNAs was induced by adding 10 µg/ml doxycycline. Samples for mRNA half-life determination were collected at 0, 2, 4, 6, 12, 18, 24, 36, 48, and 64 min for the heat shock and additional samples were collected at 90 and 120 min for the 20°C condition. At each time point, 5 ml of culture was added to an equal volume of pre-cooled methanol and stored on dry ice. RNA was extracted using the RNeasy Mini Kit.

Differential half-life (differential stability) due to codon optimality was characterized by the ratio of half-lives measured with CDSs recoded with optimal and non-optimal codons. This differential stability displays a sigmoidal dependence on the CDS length. Therefore, we fitted a logistic function to the data with the following parameters:

b and DS_Max_ represent the baseline and maximal differential stability. L_C_ and k are the threshold length and steepness parameters of differential stability, respectively. The CDS length is denoted by *x*.

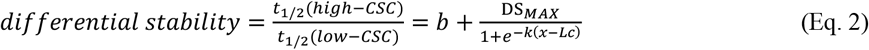

### Quantification of protein abundances with mass spectrometry

Polysome profiling was performed as described in the Polysome profiling of yeast cells section. Polysome profiling fractions were pooled based on the absorbance peaks, corresponding to the RNA complexation states (e.g. free, 40S, 60S, monosome, etc). The pooled fractions were adjusted to the same volume with nuclease free water. 1 volume of 100% trichloroacetic acid (TCA, Sigma #T0699) was added to 5 volumes of the sample to a final concentration of 17%, following which, the samples were vortexed for 10sec, the tubes wrapped with parafilm and incubated overnight at −20°C. The samples were then centrifuged at 20000g for 15min at 4°C and the supernatant was removed. 200µL of ice-cold acetone was then added gently, following which the samples were centrifuged at 20000g for 2min at 4°C. The supernatant was removed, the pellet washed by pulse vortexing with 200µL of ice-cold acetone, centrifuged at 20000g for 5min at 4°C, the supernatant again removed and the pellets air-dried. 45µL of Guanidinium-HCl buffer supplemented with 1µl of 0.75 M Chloroacetamide (CAA) was added to the samples, which were subsequently incubated at 95°C for 10min. The samples were then spun down and 135µL of 0.1M TEAB solution added, following which the pH was checked and adjusted to be higher than 7 with additional TEAB if required. 1µg of Sequencing grade Modified Trypsin (Promega #V5113) was then added to the samples, and the samples were incubated at 37°C for 12h with shaking at 300rpm. Following the trypsin digestion, the samples were acidified with 50µL of 20% trifluoroacetic acid (TFA, Sigma #T6508) and anionic exchange carried out using the Tecan Resolvex A200 automatic positive pressure processor on C18 columns. The samples were then transferred to separate 2mL tubes and dried under vacuum. Dried peptides were resuspended in 0.1% aqueous formic acid and subjected to LC– MS/MS analysis using an Orbitrap Eclipse Tribrid Mass Spectrometer fitted with an Ultimate 3000 nano-LC (both Thermo Fisher Scientific) and a custom-made column heater set to 60°C. Peptides were resolved using a RP-HPLC column (75μm × 30cm) packed in-house with C18 resin (ReproSil-Pur C18–AQ, 1.9 μm resin; Dr. Maisch GmbH) at a flow rate of 0.3 μLmin-1. The following gradient was used for peptide separation: from 2% B to 12% B over 5 min to 30% B over 40 min to 50 % B over 15 min to 95% B over 2 min followed by 11 min at 95% B. Buffer A was 0.1% formic acid in water and buffer B was 80% acetonitrile, 0.1% formic acid in water. The mass spectrometer was operated in DIA mode with a cycle time of 3 seconds. MS1 scans were acquired in the Orbitrap in centroid mode at a resolution of 60,000 FWHM (at 200 m/z), a scan range from 350 to 1200 m/z, normalized AGC target set to 250 % and maximum ion injection time mode set to 50 ms. MS2 scans were acquired in the Orbitrap in centroid mode at a resolution of 15,000 FWHM (at 200 m/z), precursor mass range of 400 to 900 m/z, quadrupole isolation window of 12 m/z with 1 m/z window overlap, a defined first mass of 120 m/z, normalized AGC target set to 1000 % and a maximum injection time of 22 ms. Peptides were fragmented by HCD (Higher-energy collisional dissociation) with collision energy set to 33 % and one microscan was acquired for each spectrum. FAIMS was set to −45V for all scans. For data analysis, SpectroNaut was used in directDIA+ mode. In brief, raw files were directly searched against a reviewed yeast uniprot FASTA database. The following search parameters were set: 2 missed cleavages, Carbamidomethylation at C as fixed modification, Oxidation at M and protein N-terminal acetylation were set as variable modifications. The output was filtered to 1% q-value cutoff and iBAQ values were selected as quantitative readout^46^.

### Analysis of protein abundances in the ribosome profiling fractions

In order to characterize the protein association with the ribosomes, the average iBAQ value of the Rps and Rpl ribosomal proteins was calculated, and the iBAQ value of each protein was divided by the average ribosome iBAQ (<RPL&RPS>). Four replicate measurements were performed to calculate the mean protein-per-ribosome ratio PpR= Protein/<RPL&RPS>. PpR at a particular temperature (e.g. PpR_30C_) was calculated for each fraction containing *i* = 1–5 ribosomes, (PpR_30C,*i*_). Proteins were excluded from further analysis if they had zero peptide count detected in any of the fractions or if the coefficient of variation calculated from four replicate measurements exceeded 1.

The monosome bias at standard conditions (30°C) is the ratio of the PpR in monosome fraction to that in the tetrasome fraction.

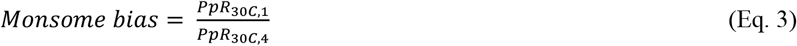

Temperature induced change in total ribosomal association (TICTRA) and temperature induced change in monosomal association (TICMA) was calculated from the PpR in the fractions with *i* ribosomes. For the TICTRA, all configurations up to *m* was included. m = 5 (pentasome+).

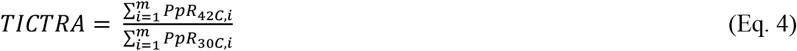

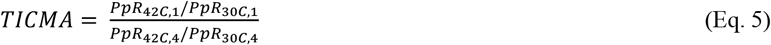

To calculate enrichment of gene ontologies, proteins that are subunits of macromolecular complexes other than ribosomes were excluded from further analysis since they may co-migrate with ribosomes in the sucrose density gradients independently. The following complexes were excluded: mitochondrial and vacuolar ATPases. These macromolecular complexes are large and/or oligomerize and migrate to densities overlapping with various cytoplasmic ribosome configurations^47,48^. Mitochondrial ribosomal proteins were excluded since they are not inhibited by cycloheximide. Mitochondria-specific proteins were excluded in general. Enrichment of gene ontology terms was performed with DAVID^49^.

### RNA metabolic labelling with 4-thiouracil (4tU)

For cell culturing, *Saccharomyces cerevisiae* cells (BY4743) were grown overnight in complete synthetic medium (Complete Supplement: Formedium #DCS0019, Yeast Nitrogen Base: Formedium #CYN0410 in 2% raffinose and 0.005% glucose).

For the 20°C set, cells were pre-cultured at 30°C for 4h and then at 20°C overnight. The overnight cultures were refreshed at an OD600 of 0.35 and grown for 4-5h until they reached the mid-log phase with a final OD600 of 0.5, then treated with 4tU.

For the 42°C set, cells were grown overnight at 30°C. The overnight cultures were refreshed at an OD600 of 0.15 and grown for 4-5h at 30°C until they reached the mid-log phase with a final OD600 of 0.5. At this point, the cells were filtered out of the media by passing through a 0.22µm membrane filter and immediately re-suspended in fresh media at 42°C. After incubation with shaking at 42°C for 30min, the culture was then treated with 4tU.

To label the mRNAs with 4tU in vivo, the cultures were treated with 4tU to a final concentration of 0.5mM and sampled. Samples were taken between 0 and 64 minutes, at intervals of power of two (0, 2, 4, 8, 16, 32 and 64min). The 0 time-point sample was taken before 4tU addition. For each time-point, 50mL of cells were collected in chilled 50% methanol. The samples were spun down for 10min at 2500g. Lysis was carried out using MPBio Teenprep™ Lysing Matrix Y tubes (MPBio # 116975050) in a FastPrep-24™ 5G bead beating grinder and lysis system. RNA was extracted with RNeasy Midi Kit (Qiagen # 75144).

For the biotinylation of labelled mRNAs, 50µg of the total RNA from each sample was mixed with an appropriate amount of spike-in RNA produced by in-vitro transcription (TranscriptAid T7: Thermo # K0441), biotinylation buffer (final concentration: 10mM HEPES [pH 7.5], 1mM EDTA) and MTSEA biotin-XX^50^ (10 µg Biotium Cat. #90066) dissolved in DMF (final concentration of DMF = 20%) in a total of 400µL. The tubes were incubated in the dark at room temperature for 2h with end-over-end rotation and the reaction was cleaned up by phenol-chloroform extraction followed by ethanol precipitation. 10% of this biotinylated RNA was kept as input.

For the enrichment of biotinylated mRNAs, 150µL of Streptavidin magnetic beads (MyOne C1 Dynabeads, Thermo: #65001) were taken per sample and separated from the liquid on a magnetic rack. Then they were washed twice with nuclease free water and then reconstituted in 100µL of nuclease free water per reaction. 100µL of the washed beads were added to 200µL of the biotinylated RNA along with 40µL of high salt wash buffer (100mM Tris-HCl [pH 7.4], 10mM EDTA, 1M NaCl, 0.1% Tween-20.) The reaction volume was made up with nuclease free water to 400µL and incubated at room temperature in the dark with end-over-end rotation for 2h.

For the elution of mRNAs from the beads, the beads were initially separated by settling on a magnetic rack, and the flow-through was discarded. The RNA-bound beads were then washed three times with 400 µL of high-salt wash buffer. The biotinylated RNA was eluted from the beads by re-suspending the beads in 100µL of freshly prepared 5% β-mercapto-ethanol (β-ME) at room temperature; incubated 15min at room temperature in the dark. The beads were separated on the magnetic rack and the supernatant collected as eluent. The beads were then again re-suspended in 100µL 5% β-ME solution at 50°C and the supernatant collected as eluent after shaking for 10min at 50°C. Both the eluents were combined, cleaned up with phenol-chloroform extraction followed by ethanol precipitation. The RNA was reconstituted in 40µL of nuclease free water supplemented with 1µL RNase inhibitor (RNasIN Plus, Promega: N2615)/100µL nuclease free water.

These eluent samples were then probed by RT-qPCR and, in addition, sent for RNASeq. rRNA depletion was down with Illumina RiboMin Gold. Libraries were generated using Illumina TruSeq Stranded Total RNA. Barcoded libraries were sequenced at single ends in Novaseq6000 (Illumina) flow cell type S1. Data processing was performed with the reference transcriptome S288C, assembly R64 (see Data processing of RNA-seq data), following which the mRNA half-lives were fitted.

### Determination of mRNA half-lives using approach to equilibrium kinetics

The half-life estimation consists of three main steps: normalization of RNA to spike-in, followed by relative normalization to intron containing nascent transcripts and completed by absolute normalization of half-lives.

For the spike-in based normalization of RNA-seq counts, the raw RNA counts at each time-point, *c = c*_*g,t*_ (where g denotes the gene and t the time-point) were divided by the spike-in RNA (Pax6) values, *f = c*_*spike-in*_, to get the spike-in normalised RNA values, *s = c* / *f*. If the number of reads at any point was less than 100, which is around 0.00125 of reads for Pax6 at 2min, the gene was discarded from calculations.

For the relative normalisation of the spike-in normalised RNA intensities, three representative nascent transcripts (i.e the introns)^29^, viz. NMD2_Nasc, ACT1_Nasc and RPL28_Nasc were used to normalise the data at each time-point in three steps.

Firstly, the average of the last two time-points (i.e 32min and 64min) of the *s*_*Nascent, t*_ series was taken where the nascent transcripts have reached the equilibrium level, *e*_*R*_ *=* mean*(s*_*Nascent, 32min*_, *s*_*Nascent, 64min*_). Then the level of the nascent RNA at each time-points were expressed as a fraction of their equilibrium levels and these values for the three nascent transcripts were averaged to give 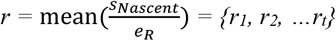.

Secondly, spike-in normalised RNA intensities of nascent mRNAs were measured by qPCR from the same RNA as the RNA-seq or from a different sample under identical conditions (explained in results section 1) to yield *m =* {*m*_*1*_, *m*_*2*_, *… m*_*t*_}. The average of the last two time-points where they have reached equilibrium was taken, i.e, *e*_*q*_ *=* mean*(m*_*Nascent,32min*_, *m*_*Nascent,64min*_). Then the level of the nascent RNA is expressed as a fraction of the equilibrium levels and these values for the three nascent transcripts were averaged to give 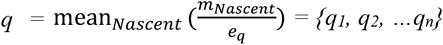.

Lastly, *u*_*t*_ *= r*_*t*_/*q*_*t*_ is calculated for each time point to rescale the spike-in normalised RNA intensities (s) at each time point as *v*_*t*_ *= s*_*t*_ */ u*_*t*_.

The data normalized to introns (*v*) were used to estimate raw half-lives by fitting the approach to equilibrium equation^7^:

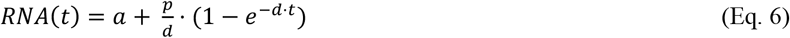

The *p* and *d* are the production and decay rate constants, respectively. The half-life of a transcript is related to *d* as 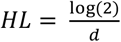. The variable *a* here refers to the non-specific binding of RNA (corresponding to the 0-time point). The R^2^ value of the fitting is also generated. The fit was carried out without any weighting. Transcripts with half-lives lower than 0.25 min or above 119 min, were discarded.

For the absolute normalisation of the half-lives, a representative set of transcripts having long and short half-lives was chosen as ‘reference’ and the half-lives of those transcripts from the metabolic labelling RNAseq were compared to the half-lives of the same genes derived by qPCR from a separate metabolic labelling experiment. Specifically, let *x =* {*x*_*1*_, *x*_*2*_, *… x*_*n*_} be the half-lives of the reference genes (1 to n) probed by qPCR and *y =* {*y*_*1*_, *y*_*2*_, *… y*_*n*_} be the half-lives of the same genes in the measured by RNASeq. *F =* mean*(y/x)* was then calculated. This *F* was then used as a scaling factor on the RNAseq metabolic labelling half-lives for an absolute scaling. *HL*_corrected_ = *HL* / *F* and thus we arrived at the normalised half-lives. Any mRNA with half-lives having a CV>0.5 (calculated from the two replicates) after the normalisation process was discarded.

## Notes

https://www.ncbi.nlm.nih.gov/geo/query/acc.cgi?acc=GSE289484

### Competing Interest Statement

The authors have declared no competing interest.

