## Supplementary Information for "Heat shock induces silent ribosomes and reorganizes mRNA turnover"

**This PDF file includes:**

Supplementary Text

Table S1

Figures S1-S7

**Other Supplementary Materials for this manuscript include the following**

Data S1 Polysome propensity and occupancy of yeast mRNAs at 42°C.

Data S2 Protein abundances in the Polysome profiling fractions

Data S3 Differential expression of yeast mRNAs due to 4-thiouracil perturbation

Data S4 Yeast mRNA half-lives estimated with metabolic labelling at 20°C and 42°C

Data S5 Polysome propensity and occupancy of macrophage mRNAs at 37°C and 42°C.

### Supplementary Text

#### Kinetic model of the cytoplasmic mRNA turnover including ribosomal association and degradation

Free mRNA (F) is produced  a rate κ most of it being cytoplasmic. Free mRNA can associate with the ribosomes, with a rate constant of $k_a$ = $k_s$· [Ribos], or degraded, with a rate constant of $\gamma_f$.

If the available ribosome concentration is reduced, for example in the presence of silent ribosomes, the association rate constant is reduced due the proportionality $k_a$ ~ [Ribos].

The mRNA bound to (i.e. translated by) Ribosomes, B, is degraded with a rate constant of $\gamma_b$.

These processes are modelled by a system of linear differential equations:

$$\frac{dF}{dt} = \kappa - \gamma_f F - k_a F \qquad \text{SEq. 1}$$

$$\frac{dB}{dt} = k_a F - \gamma_b B \qquad \text{SEq. 2}$$

In the steady-state, the mRNA occupancy (*Occup*) is the ratio of the bound mRNA (*B*) to the total mRNA (*F* + *B*).

$$Occup = \frac{B}{F+B} \qquad \text{SEq. 3}$$

Combining the bound mRNA (from SEq. 2, steady-state) and SEq. 3 yields:

$$Occup = \frac{\frac{k_a F}{\gamma_b}}{F + \frac{k_a F}{\gamma_b}} = \frac{k_a}{\gamma_b + k_a}$$

Thus, the ribosomal association rate constant ($k_a$) is a function of mRNA occupancy and mRNA degradation rate constant, $\gamma_b$.

$$k_a = \gamma_b \frac{Occup}{1 - Occup}$$

#### Supplementary Tables

Table S1: Transcript half-lives (in min) of MGC genes at 42°C (30’) treated with 4tU for 6’ (3 replicates)

| Gene | DMSO Half-life | SEM DMSO | 4tU Half-life | SEM 4tU | FC Half-life | SEM FC |
|---|---|---|---|---|---|---|
| *OLA1* | 5.12 | 0.09 | 10.58 | 0.81 | 2.07 | 0.18 |
| *RFS1* | 3.91 | 0.58 | 7.25 | 1.71 | 1.85 | 0.26 |
| *NRG2* | 1.56 | 0.16 | 4.43 | 0.83 | 2.82 | 0.32 |
| *GAL80* | 2.15 | 0.20 | 6.08 | 0.80 | 2.81 | 0.21 |
| *IMD4* | 3.14 | 0.32 | 7.81 | 1.52 | 2.43 | 0.25 |
| *TEM1* | 0.88 | 0.09 | 3.40 | 0.38 | 3.97 | 0.63 |
| *DAK1* | 4.95 | 0.25 | 4.42 | 0.53 | 0.89 | 0.07 |
| *FPR3* | 4.27 | 0.35 | 8.14 | 0.64 | 1.91 | 0.08 |
| *RAD10* | 0.97 | 0.12 | 3.81 | 0.79 | 3.85 | 0.35 |
| *PLM39* | 0.57 | 0.06 | 2.38 | 0.70 | 4.06 | 0.74 |
| *DAT1* | 1.69 | 0.09 | 5.89 | 0.24 | 3.49 | 0.10 |
| *NUP188* | 2.22 | 0.10 | 4.35 | 0.22 | 1.97 | 0.08 |
| *SML1* | 2.81 | 0.32 | 3.99 | 0.37 | 1.44 | 0.11 |
| *MIS1* | 2.58 | 0.17 | 5.60 | 0.68 | 2.16 | 0.14 |

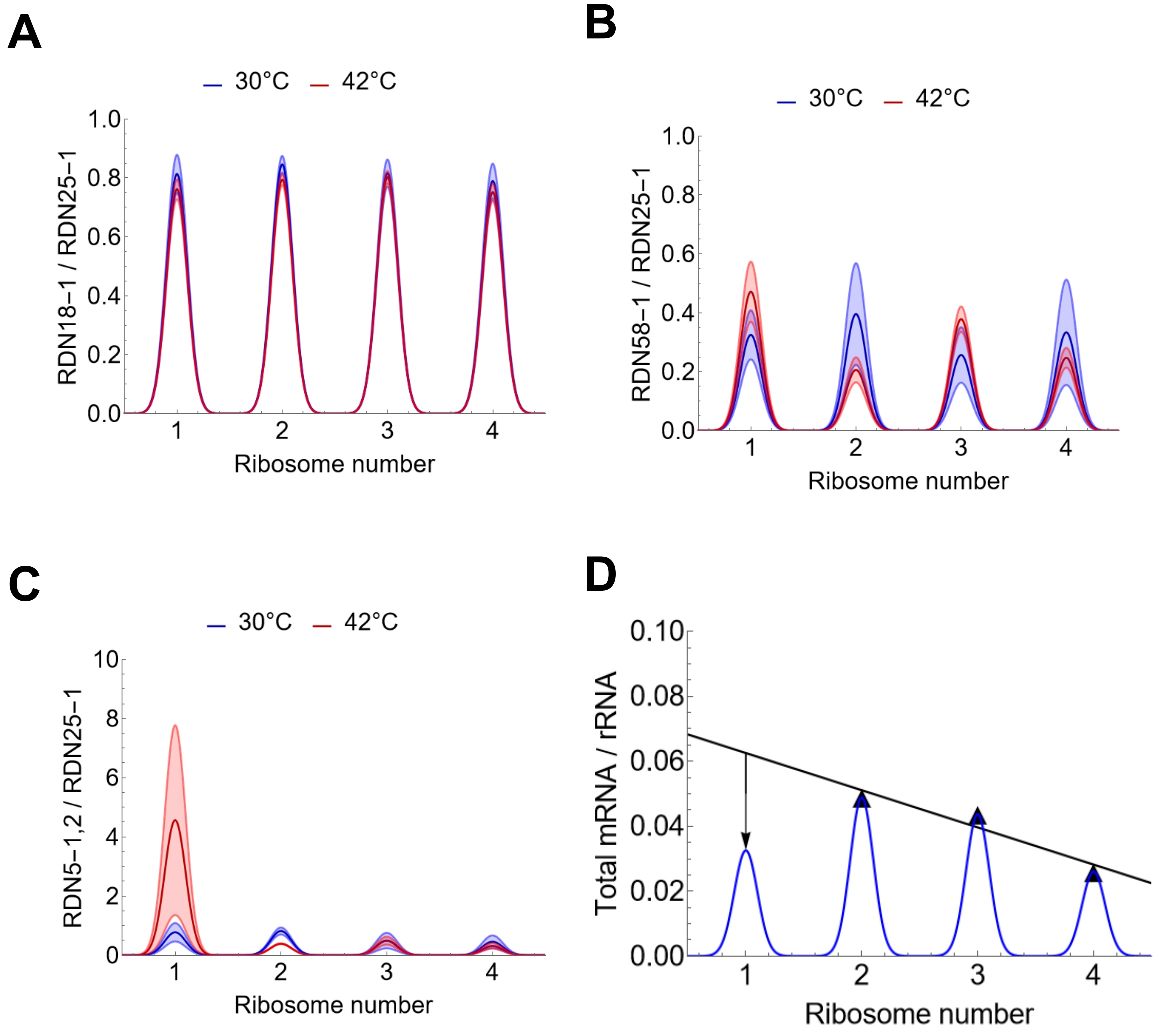


**Figure S1. Measurement of rRNAs and estimation of silent ribosome proportions. (A-C)** Consistency of the 18S / 25S, 5.8S / 25S and 5S / 25S rRNA ratios measured with RNA-seq in the non-depleted RNA samples isolated from the polysome profiling fractions with the indicated ribosome numbers. The thick dark and the thin light lines represent **(D)** Linear regression of ∑mRNA/∑rRNA values measured at 30°C for configurations with $N$ = 2, 3, and 4 ribosomes (black triangles). The deviation of the measured value at the monosome ($N$ = 1) from the regression line is indicated by an arrow.

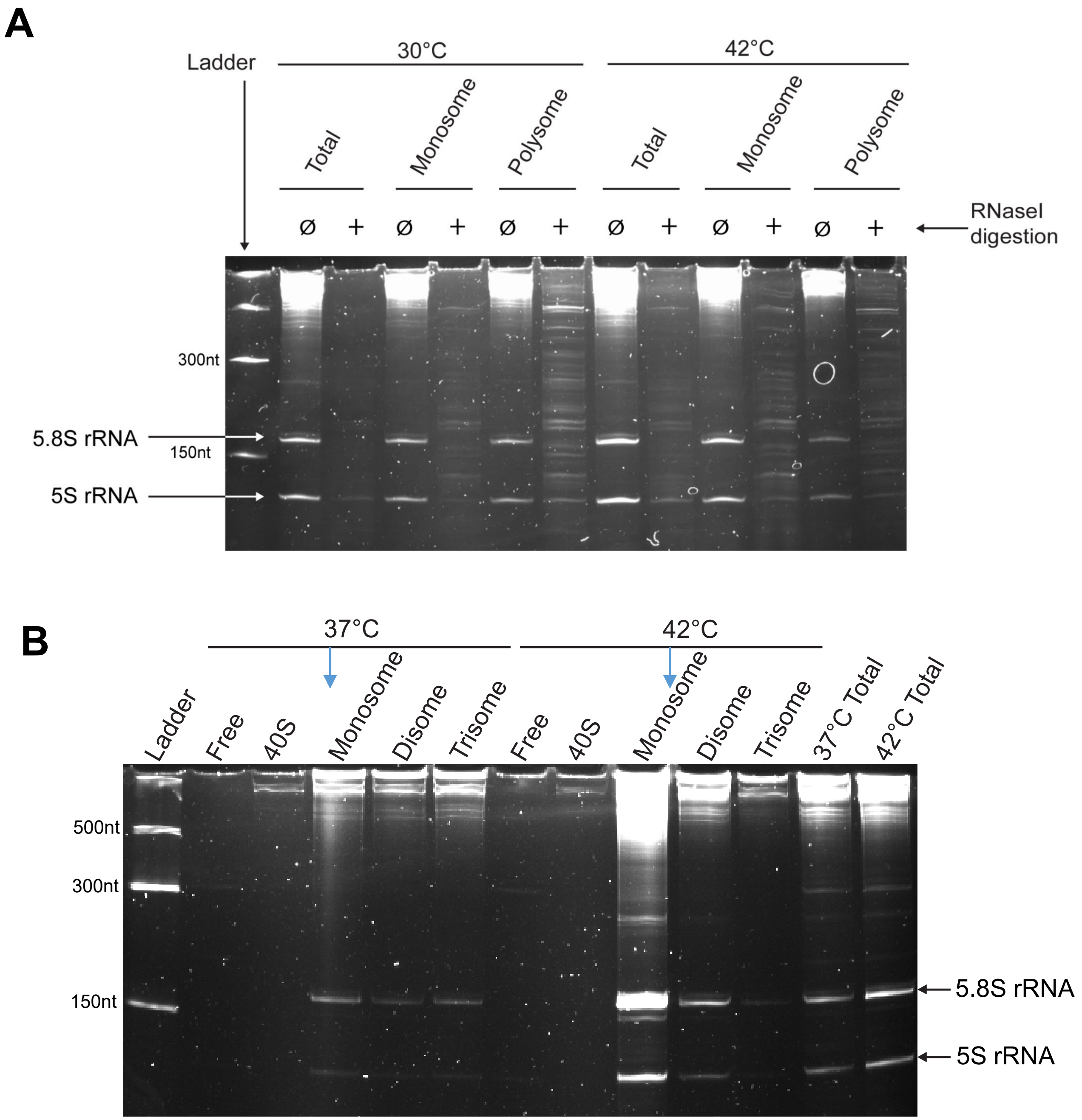


**Figure S2. rRNA intensities measured with PAGE.** Yeast (A) or J774A.1 macrophage (B) total RNA as well as RNA extracted from monosomal and different polysomal fractions of the polysome profile were separated on an 8% polyacrylamide gel with 8M urea. Band intensities were quantified using FIJI, with background subtraction performed using an empty region of the gel. Both 5.8S and 5S rRNAs show an increase in the monosomal fraction after heat shock. The following fold change in RNA intensity due to heat shock [FC] were measured. **(A)** FC(42°C/30°C) in yeast, FC(5S rRNA) = 1.45; FC(5.8S rRNA) = 1.40,). There is no anomalous amplification of 5S rRNA (FC ratio [FC(5S)/FC(5.8S)] = 1.04). **(B)** FC(42°C/37°C) in macrophage, FC(5S rRNA) = 3.44; FC(5.8S rRNA) = 3.87,). There is no anomalous amplification of 5S rRNA (FC ratio [FC(5S)/FC(5.8S)] = 0.89).

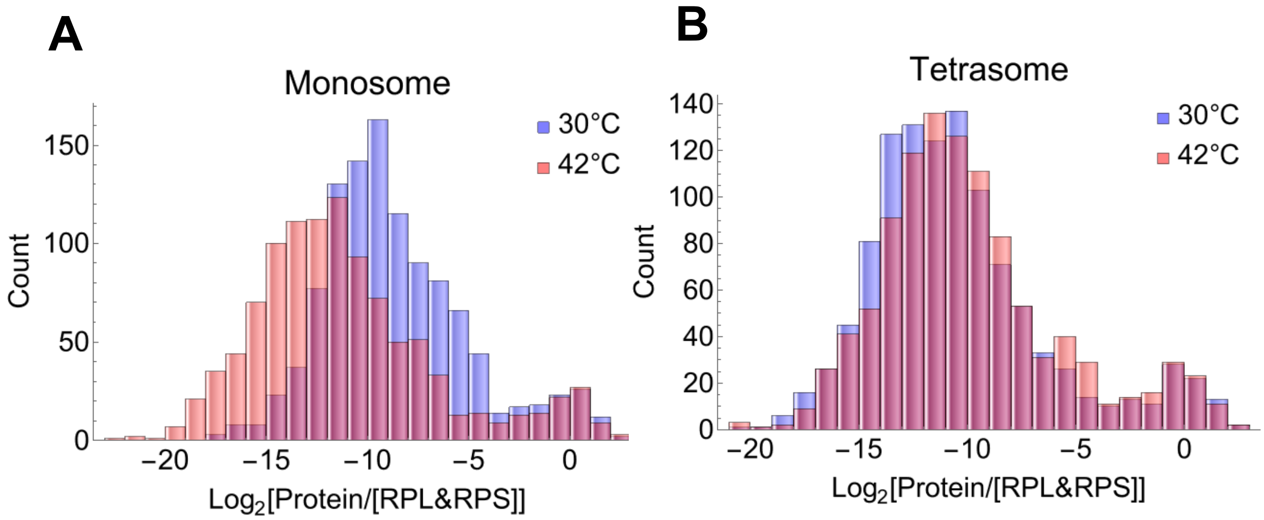


**Figure S3. Distribution of protein abundances in the monosome and tetrasome fractions. (A)** Comparison of proteins in the monosome fractions. The median normalized abundance of the top 100 proteins is 0.70 at 30°C and 0.66 at 42°C (P-value = 0.54. Mann-Whitney test). The median abundance of all other proteins is 0.00132 at 30°C and 0.00017 at 42°C (P-value = $1.48\cdot10^{-61}$. Mann-Whitney test). Among the top 100 proteins at 30°C, 81 are ribosomal proteins, including 45 from the large subunit (RPL), 33 from the small subunit (RPS), and 3 from the stalk (RPP). These results indicate that while the smaller peaks, composed primarily of ribosomal proteins, are comparable between conditions, the larger peak of ribosome-associated proteins shows a 7.7-fold decrease in abundance at 42°C. **(B)** Comparison of proteins in the tetrasome fractions. The median normalized abundance of the top 100 proteins is 0.70 at 30°C and 0.69 at 42°C (P = 0.71, Mann-Whitney test). The median abundance of all other proteins is 0.00033 at 30°C and 0.00043 at 42°C (P = 0.007, Mann-Whitney test). Among the top 100 proteins at 30°C, 81 are ribosomal proteins, including 46 from the large subunit (RPL), 32 from the small subunit (RPS), and 3 from the stalk (RPP). These results suggest that the smaller peaks, containing mostly ribosomal proteins, remain consistent between conditions, while the larger peak of ribosome-associated proteins shows a modest 1.3-fold increase in abundance at 42°C.

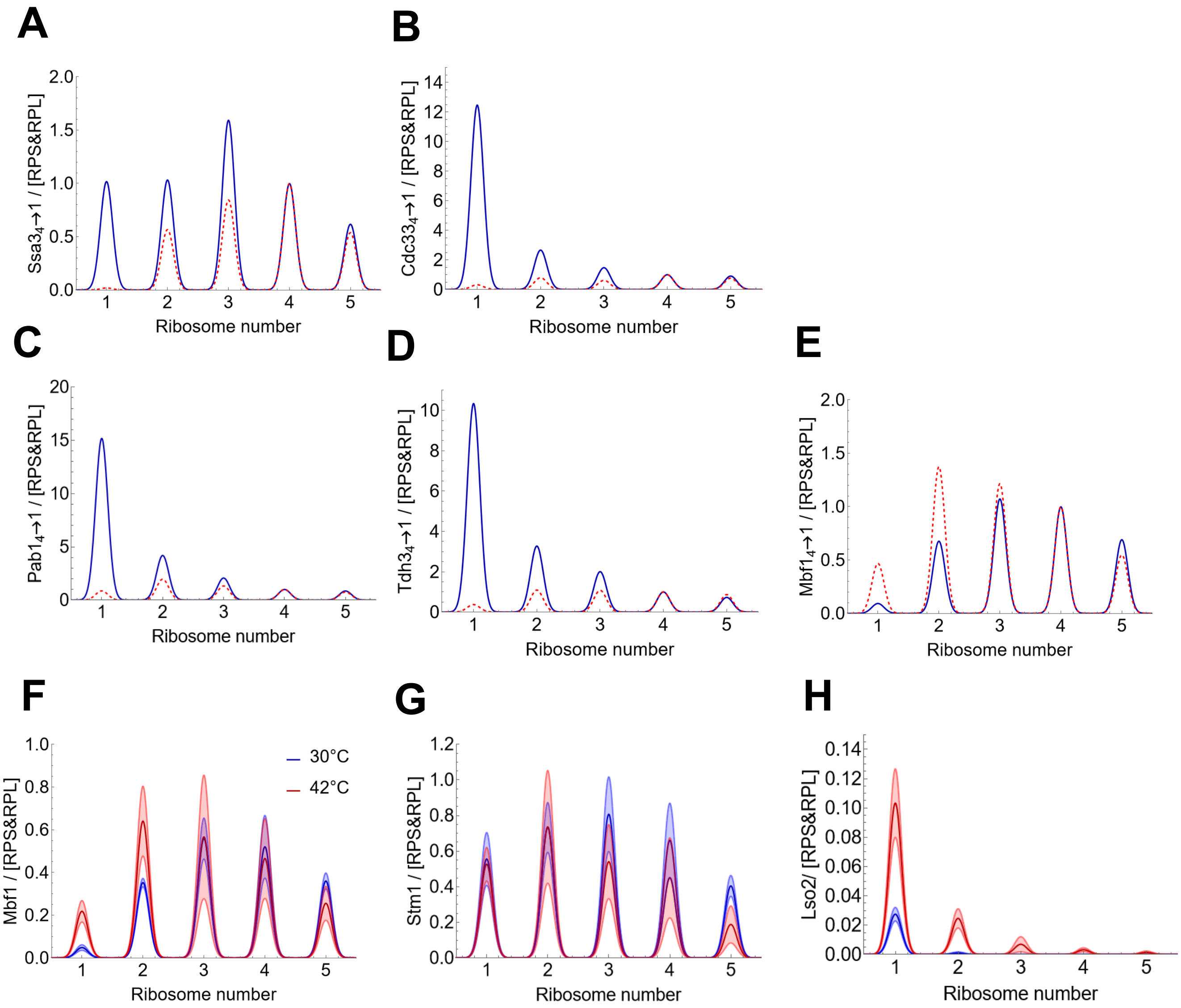


**Figure S4. Proteomic analysis of the polysome profiling fractions. (A-E)** Protein abundances normalized relative to the tetrasome level. Protein/<RPL&RPS> (PpR) ratios measured at both 30°C (blue full lines) and 42°C (red dashed line) were set to 1 in the tetrasome fraction. **(F-H)** The dark and light lines show the mean and standard deviation (n = 4) of the protein abundances of the indicated proteins normalized to the ribosomal average in the indicated fractions of the polysome profiling.

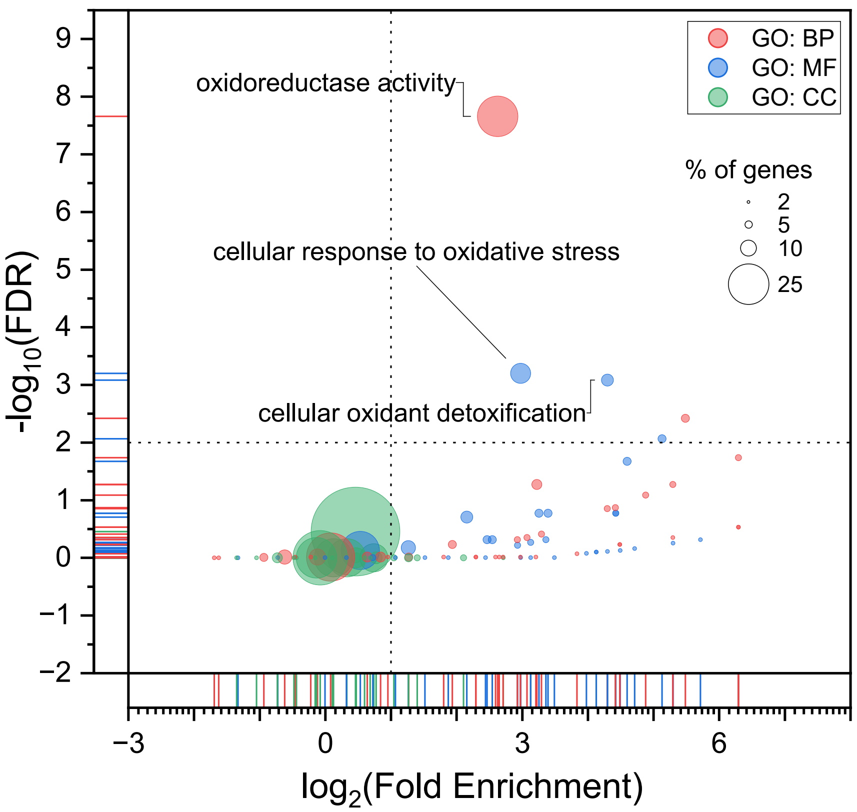


**Figure S5. Gene Ontology (GO) analysis of the genes differentially expressed upon 4tU treatment at 30°C and 42°C.** A total of n=82 genes are differentially expressed under both conditions (adjusted p-value < 0.05). Enrichment analysis of these genes was performed for three GO categories: biological process (BP, red), molecular function (MF, blue) and cellular component (CC, green). Lowest false discovery rates (FDR) were found for GO terms related to redox activity and cellular stress. Bubble sizes represent the percentage of genes associated with particular GO term among the 82 differentially expressed genes.

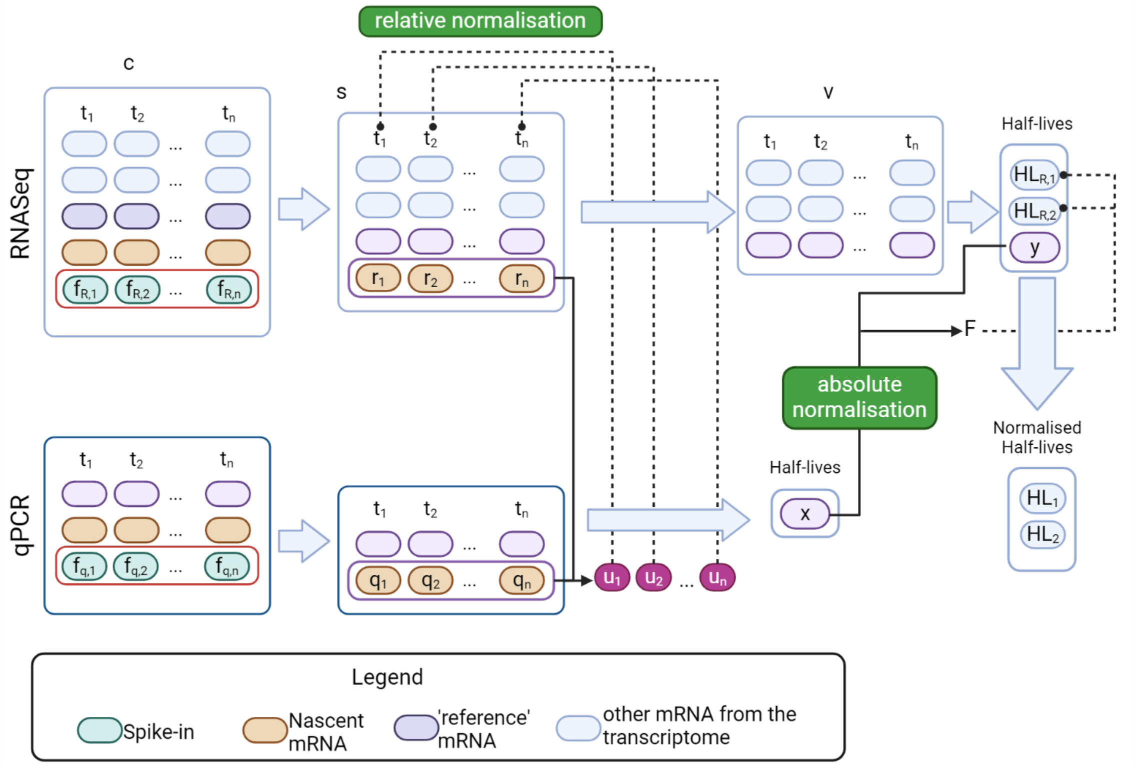


**Figure S6**. **Flowchart depicting steps to determine mRNA half-lives from metabolic labelling.** The raw RNA counts (*c)* are first converted to spike-in normalised RNA values (*s*), followed by relative normalization yielding RNA intensities (*v)* (see Methods for further details). $t_i$ refers to the time-point *i*. *f* denotes the raw RNA count or intensity of spike-in RNAs measured by RNAseq ($f_R$) or by qPCR ($f_q$). $r_i$ and $q_i$ represent nascent transcript levels expressed as fractions of their equilibrium value at time point *i* measured by RNAseq and qPCR respectively. *x* and *y* refer to the half-lives of reference mRNAs measured by qPCR and RNAseq, respectively. *F* is the absolute normalisation factor.

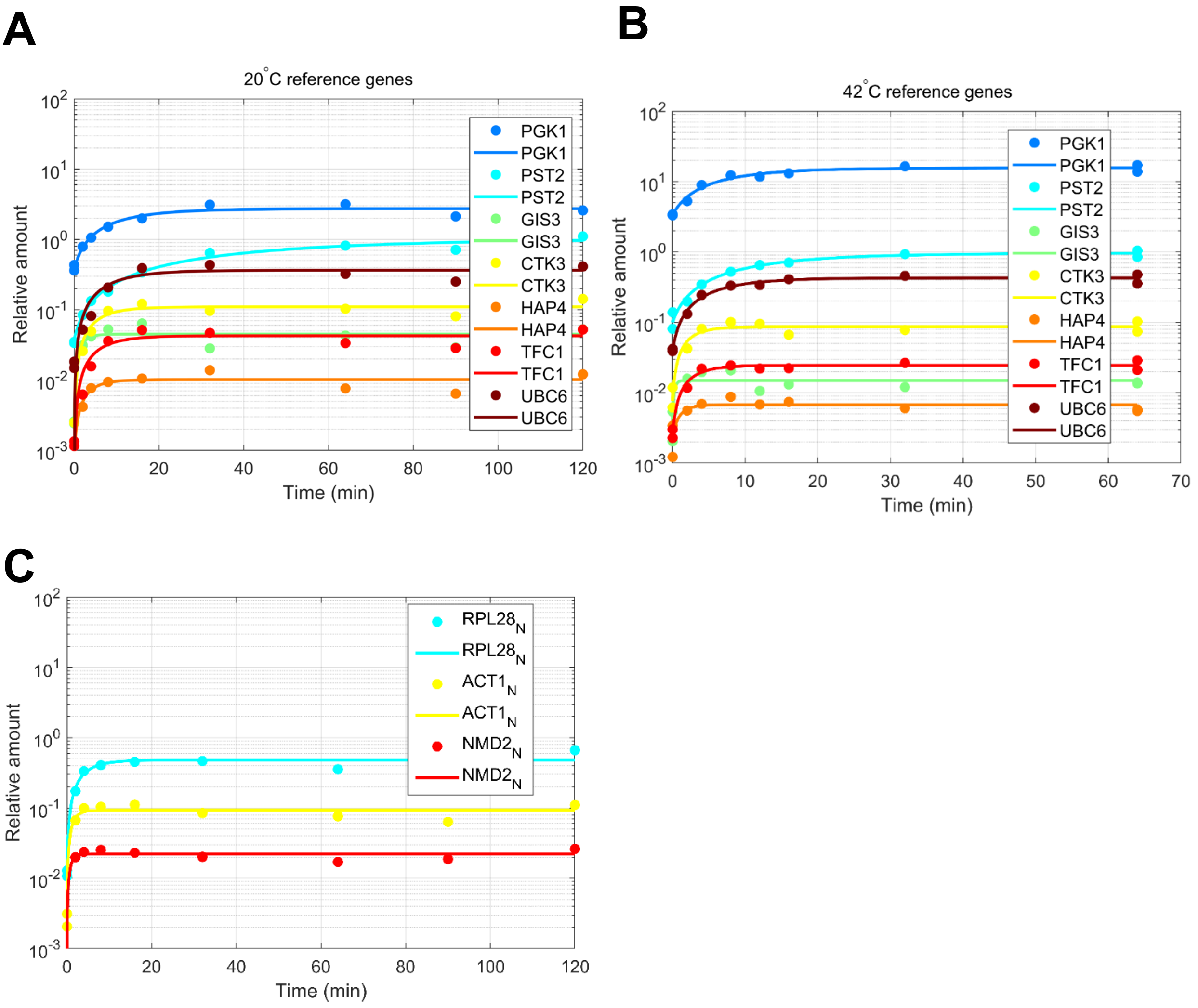


**Fig. S7. Time series of labelled reference and nascent unspliced mRNAs.** mRNAs eluted by β-mercaptoethanol were quantified by qPCR. Time series of the 'reference' genes from the absolute normalization samples fitted with Equation 6 (Methods) (**A, B**) qPCR based RNA abundances were normalized by spike-in and nascent RNAs ($v$ set). The following half-lives were fitted at 20°C and 42°C respectively: *PGK1*: 7.70 and 5.76 min; *PST2*: 29.77 and 9.03 min; *GIS3*: 1.13 and 0.20 min; *CTK3*: 3.63 and 1.56 min; *HAP4*: 2.39 and 0.98 min; *TFC1*: 4.01 and 1.81 min; and *UBC6*: 5.77 and 4.42 min. (**C**) Time series of three nascent genes (unspliced) fitted with Equation 6 (Methods) following metabolic at 20°C. qPCR based RNA abundances were normalized by spike-in ($q_i$ ). Due to the fast turnover of nascent mRNAs as a result of rapid splicing, their levels quickly reach saturation.
